# Mapping the dynamic cell surface interactome of high-density lipoprotein reveals Aminopeptidase N as modulator of its endothelial uptake

**DOI:** 10.1101/2023.01.03.522574

**Authors:** Kathrin Frey, Lucia Rohrer, Anton Potapenko, Sandra Goetze, Arnold von Eckardstein, Bernd Wollscheid

## Abstract

Heterogeneous high-density lipoprotein (HDL) particles, which can contain hundreds of proteins, affect human health and disease through dynamic molecular interactions with cell surface proteins. How HDL mediates its long-range signaling functions and interactions with various cell types is largely unknown. Due to the complexity of HDL, we hypothesize that multiple receptors engage with HDL particles resulting in condition-dependent receptor-HDL interaction clusters at the cell surface. Here we used the mass spectrometry-based and light-controlled proximity labeling strategy LUX-MS in a discovery-driven manner to decode HDL-receptor interactions. Surfaceome nanoscale organization analysis of hepatocytes and endothelial cells using LUX-MS revealed that the previously known HDL-binding protein scavenger receptor SCRB1 is embedded in a cell surface protein community, which we term HDL synapse. Modulating the endothelial HDL synapse, composed of 60 proteins, by silencing individual members showed that the HDL synapse can be assembled in the absence of SCRB1 and that the members are interlinked. The aminopeptidase AMPN (also known as CD13) was identified as an HDL synapse member that directly influences HDL uptake into the primary human aortic endothelial cells (HAECs). Our data indicate that preformed cell surface residing protein complexes modulate HDL function and suggest new theragnostic opportunities.

## Introduction

The cell surface is not only a means for the compartmentalization of a cell but also serves as a gateway to the exterior and as a signaling mediator between extracellular stimuli and intracellular responses (1). The surfaceome is a dynamic organization of functional clusters of proteins that interact with extrinsic ligands (2). It has become clear that the nanoscale organization of receptors is a critical mediator of molecular function in health and disease. For instance, virus particles bind simultaneously to multiple receptors increasing their affinity for the host cell and likely induce signaling pathways by receptor clustering through many-to-many interactions (3).

The heterogenous high-density lipoprotein (HDL) particles engage with surfaces of different cells in systemic manner, contributing to the protection of the human body from many diseases (4). HDL particles contain hundreds of proteins, different lipid species, and microRNAs (4, 5). Structural heterogeneity implies functional heterogeneity: In addition to mediating reverse cholesterol transport, HDL particles trigger a wide range of cytoprotective signaling events including the induction of vasoprotective effects by nitric oxide stimulation, maintenance of endothelial barrier integrity, and the mediation of anti-apoptotic, anti-inflammatory, and antithrombotic effects (6). In 1996, Acton and colleagues reported that hepatocytes bind HDL particles with high affinity through the scavenger receptor SCRB1 to mediate selective uptake of cholesterol without internalizing the entire particle (7). SCRB1 is a widely expressed, multiligand receptor that serves as a multifunctional platform involved in various HDL-triggered signaling events as well as HDL holo-particle uptake into some non-hepatic cells including endothelial cells (8, 9). For many functions of HDL, SCRB1 is neither necessary nor sufficient. In endothelial cells, SCRB1, sphingosine-1-phosphate receptors, and the ectopic beta-ATPase were all found to limit the ability of HDL particles to promote nitric oxide production (10). This suggests that SCRB1 either interacts with or is functionally redundant with other receptors. A lack of knowledge regarding the spatial nanoscale organization of the HDL interactome, which ultimately manifests HDL functionality, has hampered the uncovering of the structure-function relationship of HDL particles.

Due to the complexity of the HDL particle, we hypothesized that multiple receptors participate in HDL binding, uptake, and signaling, resulting in condition-dependent HDL synapses at the cell surface. To understand such complex molecular interaction networks, experimental approaches with spatial resolution are required. The mass spectrometry-based, light-controlled proximity labeling strategy LUX-MS enables the characterization of complex ligand-receptor interactions (11): Receptors proximal to a cell surface-bound ligand are tagged in a light-dependent manner and can subsequently be identified by mass spectrometry. Moreover, via automated Cell Surface Capturing (auto-CSC), cell surfaceomes can be characterized qualitatively and quantitatively in a holistic manner (12).

To decipher the *bona fide* HDL synapses of endothelial cells, we utilized the LUX-MS platform to directly identify HDL-residing proteins and their receptor neighborhoods at the cell surface. We created an inventory of functional HDL-receptor neighborhoods on endothelial cells and compared it to hepatic HDL synapses. By assessing the quantitative endothelial surfaceome remodeling upon intrinsic perturbation of the HDL receptor nanoscale organization by silencing of individual HDL receptor candidates, we found that HDL receptor candidates are interlinked. These receptors include the aminopeptidase AMPN (also known as CD13). In proof-of-principle experiments, we validated AMPN as an important component of the HDL synapse that contributes to HDL uptake into primary human endothelial cells.

## Results

### The endothelial HDL synapse includes the SCRB1 receptor environment but is also assembled in the absence of SCRB1

To map the HDL synapse, we applied the LUX-MS strategy to human endothelial somatic hybrid (EA.hy926) cells using HDL-singlet oxygen generator (SOG) constructs (**Figure 1A**). Flow cytometry revealed the biotinylation rate at the cell surface of EA.hy926 cells as a function of illumination time (**Figure 1B**). Maximum biotinylation was observed at 15 min, and we therefore chose this illumination time for all subsequent LUX-MS experiments. To validate our LUX-MS strategy, we qualitatively compared anti-SCRB1 antibody-based and HDL-based LUX-MS snapshots. As the low abundance of SCRB1 on wild-type EA.hy926 cells did not permit an anti-SCRB1 antibody-based LUX-MS, we used SCRB1 variant 1 (V1) overexpressing cells for this comparison (**Suppl. Figure 1A and B**). In the experiment using anti-SCRB1 antibody, the ligand (anti-SCRB1 mouse IgG) and the target protein SCRB1 were enriched compared to the control condition as were 49 other proteins (**Figure 1C, Suppl. Figure 2, Suppl. Table 1**). Of these 49 proteins, 38 were also enriched in the HDL-based LUX-MS experiment (**Figure 1D**). However, the HDL-based LUX-MS snapshot contained 63 additional proteins, including the main protein of HDL, the apolipoprotein APOA1. The large coverage of the SCRB1 neighborhood in the HDL-based LUX-MS experiment showed that our strategy captures the HDL synapse. These results also indicated that HDL interacts with proteins beyond SCRB1 and its direct environment on endothelial cells. In line with this observation, while HDL binding to EA.hy926 cells overexpressing SCRB1 was enhanced relative to wild-type cells, the interaction with HDL was not entirely prevented by reduction of the cell surface abundance of SCRB1 to less than 5% compared to control using the RNA interference strategy (**Suppl. Figure 1C-D and 3)**. In these SCRB1-deficient cells, an HDL-guided LUX-MS experiment recovered almost 80% of the proteins that were identified in cells that overexpress SCRB1 (**Figure 1E and F, Suppl. Table 1**). This indicates that the SCRB1 neighborhood plays an important role in HDL binding even at very low abundance of SCRB1. These results indicate that HDL particles bind to the SCRB1 environment through interactions that are dependent on and independent of SCRB1.

**Figure 1:**
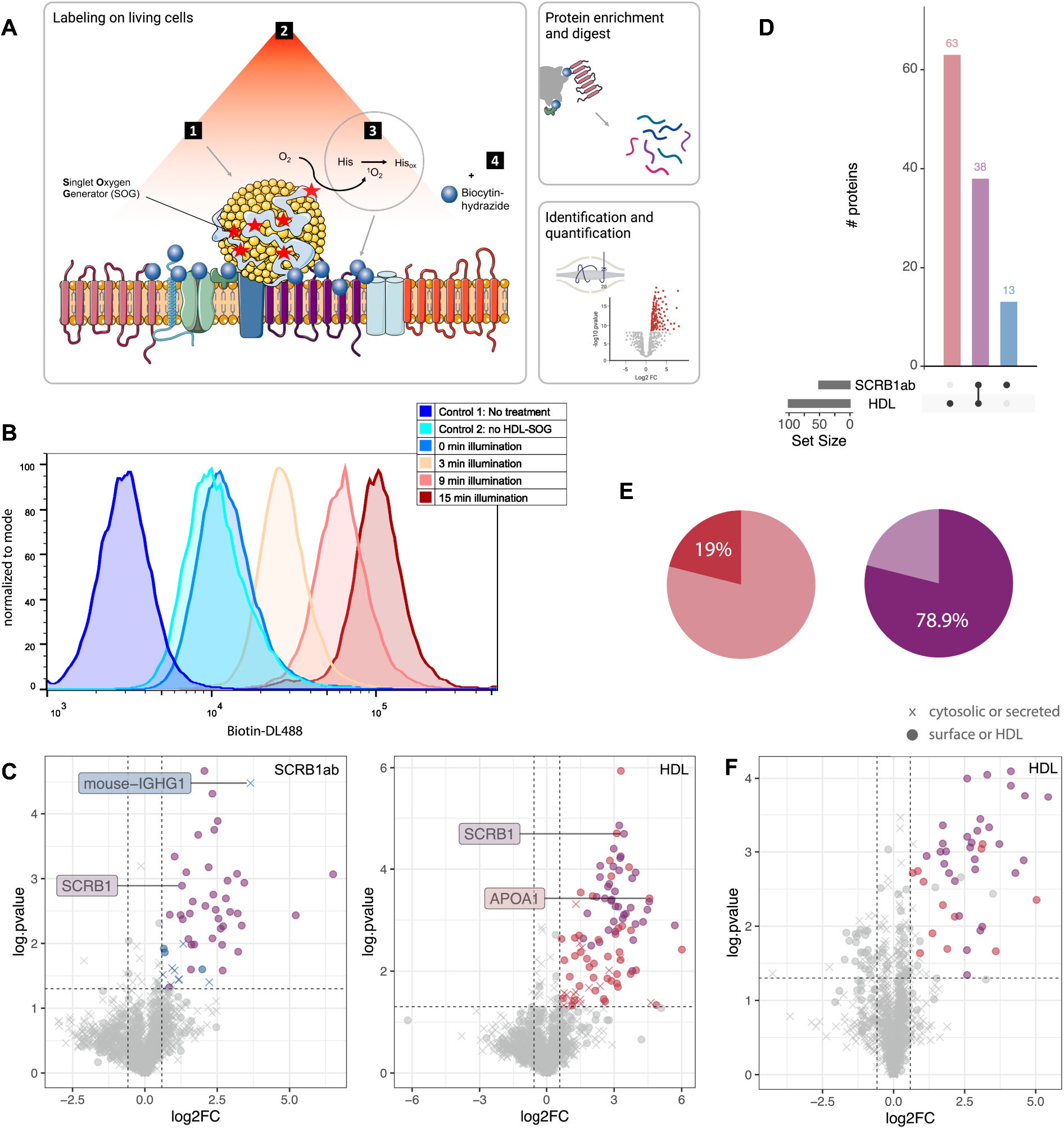
Characterization of HDL synapses using the LUX-MS technology. **A)** Schematic illustration of the LUX-MS workflow: (1) The SOG-decorated ligand (here HDL) is bound to the living cell. (2) Upon illumination, singlet oxygen is generated. (3) Singlet oxygen modifies proximal amino acids. (4) Modified amino acids are subsequently biotinylated. Following these labeling steps, the cells are lysed, biotinylated proteins enriched and proteolytically released. The peptides are then identified by liquid chromatography tandem mass spectroscopy (LC-MS/MS) and quantified against the control (no SOG). **B)** Flow cytometry-based assessment of biotinylation of wild-type EA.hy926 cells during the LUX reaction using HDL as a ligand with different illumination times. **C)** Volcano plots of LUX-MS experiments using an anti-SCRB1 antibody (left) and HDL (right) as ligands on SCRB1 V1-overexpressing EA.hy926 cells. Proteins enriched in the anti-SCRB1 antibody-based LUX-MS experiment are in blue. Proteins enriched in the HDL-based LUX-MS experiment are in red. Proteins enriched in both experiments are in purple. **D)** Numbers of proteins identified in the anti-SCRB1 antibody-based (red) and HDL-based (blue) LUX-MS experiments and the number of overlapping proteins (purple). **E)** Fraction of proteins enriched in HDL-based LUX-MS experiment on EA.hy926 cells in which SCRB1 was silenced (dark red, dark purple) compared to enriched proteins on SCRB1 V1-overexpressing cells (colors as in panel C). **F)** Results of the HDL-based LUX-MS experiment on EA.hy926 cells in which SCRB1 was silenced displayed as a volcano plot using the same color code as in panel C. Significance was determined using MSstats (fold change (FC) > 1.5 and p-value < 0.05).

### At least 60 proteins define the endothelial HDL synapse

LUX-MS labels both cell-derived and HDL-derived proteins. The HDL-residing proteins might contribute to the more complex enrichment pattern observed in the HDL-guided LUX-MS experiment compared to the anti-SCRB1 antibody-guided LUX-MS experiment. HDL particles are composed of highly diverse and variable protein cargos, including cellular proteins that probably associate with HDL either during cellular trafficking or after shedding (5). To discriminate between HDL- and cell-derived proteins and to investigate the variability within the HDL synapse, we qualitatively analyzed and compared four different HDL-based LUX-MS snapshots of wild-type EA.hy926 cells (**Suppl. Figure 4, Suppl. Table 2 and 3**). For half of these snapshots, we grew wild-type EA.hy926 cells in SILAC medium containing the heavy isotope-labeled arginine and lysine to enable the discrimination between HDL-associated and EA.hy926-derived proteins. In these experiments, HDL-derived proteins and EA.hy926-derived proteins were identified as light and heavy peptides, respectively. Of the 22 proteins identified with light peptides, 11 were identified only through detection of light peptides and hence were derived from the HDL particle, whereas 11 proteins were identified by detection of both light and heavy peptides (**Figure 2A)**. This former group of proteins are considered to be part of both HDL and the cell surface of EA.hy926 cells. By comparing our findings with the surface proteome of wild-type EA.hy926 cells and/or an inventory of the HDL proteome ((5, 13), and http://homepages.uc.edu/~davidswm/HDLproteome.html, March 2021), nine proteins could be validated to be part of the HDL proteome and/ or the EA.hy926 surfaceome, respectively. Thirteen proteins had not previously been associated with HDL particles.

**Figure 2:**
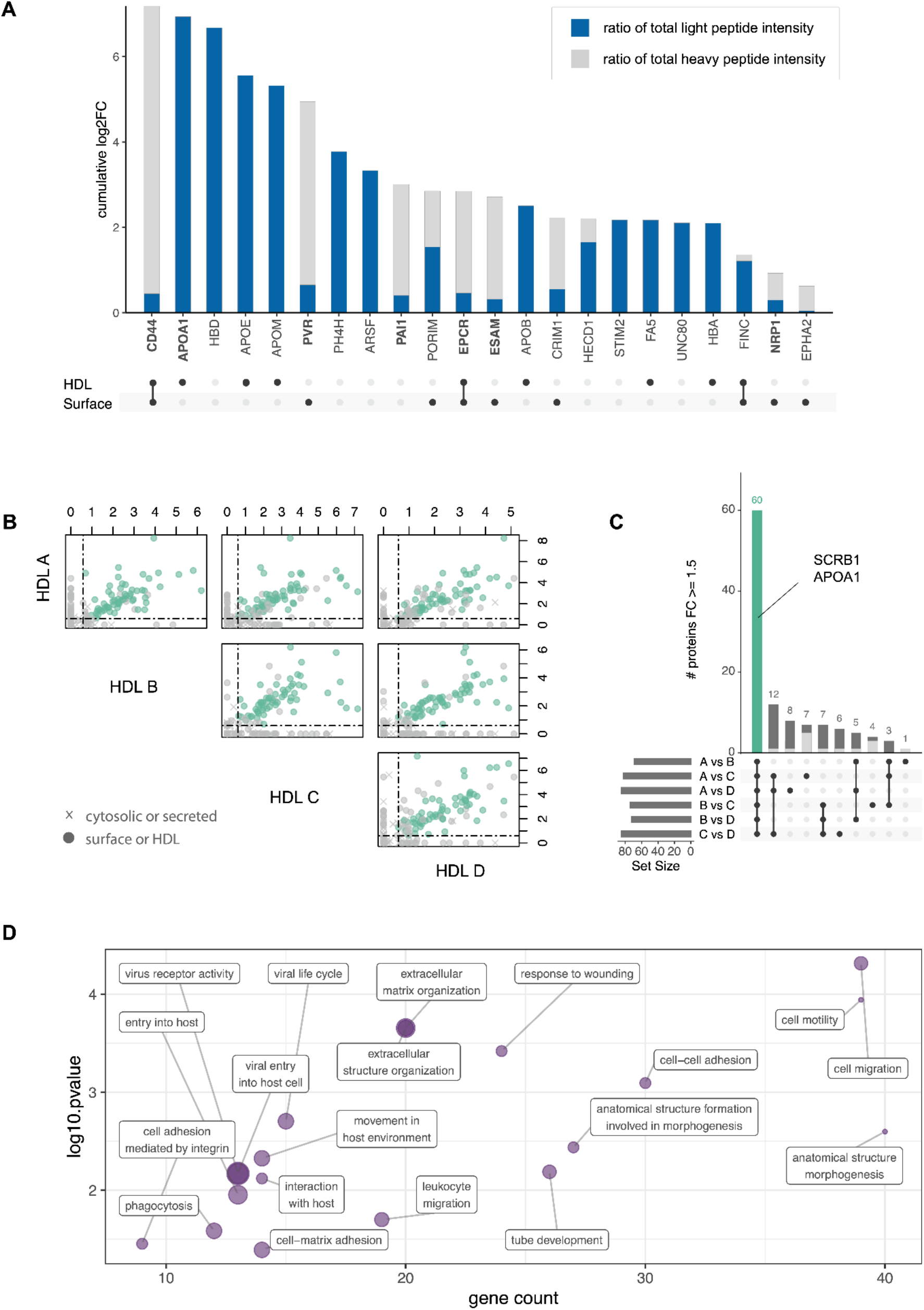
Decoding the HDL-derived and cell-derived HDL synapse compositions across four independent experiments. **A)** Cumulative log2(FC) for indicated proteins in the four HDL-based LUX-MS experiments relative to corresponding controls. In each bar, the blue region indicates the ratio of light peptide intensities to heavy peptides (mean of two experiments). Proteins with a FC > 1.5 in all four experiments are highlighted in bold. Dots below the graph indicate whether the protein is part of the Surfy or the HDL proteome list. **B)** Correlation plot with log2(FC) of the enriched proteins in the four independent LUX-MS experiments with four different HDL isolates (HDL A - D). Green points indicate FC > 1.5 in all four experiments. **(C)** Number of proteins that reached a FC threshold > 1.5 in all four experiments. Dark grey indicates surface or HDL annotation for the protein and light grey indicates a cytosolic protein. Green bar indicates the 60 core candidates; all are on the Surfy or the HDL proteome list. **D)** Gene ontology analysis of the 60 core candidates compared to the endothelial cell surfaceome. Displayed are all significantly enriched terms (p value corrected with Bonferroni step down < 0.05). The size of the dots inversely corresponds to the mean gene ontology term level.

Next, we assessed the variance of the HDL- and cell-derived components within the HDL synapses across four independent HDL-based LUX-MS snapshots. The qualitative and quantitative proteomes of HDL particles used for these HDL synapse snapshots varied between isolates, but 60 proteins were enriched in all four snapshots, indicating that the HDL composition only slightly influences the qualitative composition of the HDL synapse (**Figure 2B and C, Suppl. Figure 2, Suppl. Figure 4, Suppl. Table 1**). Of the 60 proteins enriched in all snapshots, 60% were also detected in an HDL-based LUX-MS experiment on primary human aortic endothelial cells (HAECs) (**Suppl. Figure 5, Suppl. Table 1**). Despite the low abundance of SCRB1 on wildtype EA.hy926 cells, SCRB1 was among the 60 proteins detected in these cells that we, from now on, refer to as the HDL synapse core candidates. APOA1 was the only solely HDL-derived protein within this HDL synapse core candidate list. Although heavy peptides were most prominent, light peptides were also detected for CD44, poliovirus receptor PVR, plasminogen activator inhibitor PAI1, endothelial protein C receptor EPCR, endothelial cell-selective adhesion molecule ESAM, and neuropilin NRP1 (**Figure 2A)**. All other HDL synapse core candidates were annotated as surface receptors. Three candidates have transporter activities: sodium/potassium-transporting ATPase subunits AT1B1 and AT1B3 and cell-surface antigen heavy chain 4F2. Eight proteins have enzymatic activities including tyrosine protein kinase receptors UFO and TIE1 and the receptor-type tyrosine-protein phosphatases PTPRB and PTPRF. The remaining 21 proteins with receptor function include two scavenger receptor family members, SCRB1 and SREC (14). We performed a gene ontology biological processes analysis to gain a broad overview of the functionalities of the 60 candidates (**Figure 2D, Suppl. Table 4**). Most receptors were associated with anatomical structure morphogenesis, cell migration, and motility. Six out of 19 enriched terms are related to virus interactions with the host. The main drivers behind enrichment in these six terms are SCRB1, PVR, NECT2, AMPN, UFO, DAG1, the low-density lipoprotein receptor LDLR, and six integrin family proteins.

### Surfaceome analysis of the dynamic interplay of the endothelial HDL synapse

We next set out to study the dynamic interplay of the 60 HDL synapse candidates. The HDL-guided LUX-MS experiments on EA.hy926 cells revealed that the endothelial HDL synapse remains largely unaltered upon silencing or overexpression of SCRB1 (**Figure 1E**). This suggests that the cell surface proteins that interact with HDL particles adapt to such perturbations to enable HDL-relevant functionalities. To study remodeling of the proteome at the cell surface upon longterm perturbation, we used the auto-CSC platform, which enables the identification and quantification of *N*-glycosylated cell surface receptors (12). We stably silenced the HDL core candidates SCRB1, adhesion molecule PECA1, thrombin modulator TRBM, and tyrosine-protein phosphatase substrate SHPS1 individually in EA.hy926 cells by expression of shRNAs from viral vectors (**Suppl. Figure 6**). Furthermore, we silenced expression of NECT2, which is a potential SCRB1-independent HDL interactor because it was enriched by the HDL-guided LUX-MS experiment but not in the anti-SCRB1-guided LUX-MS experiment. NECT2 is a receptor for virus entry (15) and has been shown to be cholesterol responsive and to influence atherosclerosis development by affecting leukocyte migration (16). We observed quantitative surfaceome reorganizations in the cell lines deficient in HDL core candidates compared to the control cell surfaceome treated with a control plasmid (**Suppl. Figure 7A-E, Suppl. Table 5 and 6**). Lack of SHPS1 resulted in the most extensive surfaceome reorganization with 20 upregulated and 17 downregulated proteins. For instance, adhesion molecule ICAM1 was upregulated upon SHPS1 silencing. ICAM1 expression was previously shown to be suppressed by SCRB1 dependent and independent mechanisms (17); the SCRB1 independent mechanism involves HDL-associated sphingosine-1-phosphate (18). It is noteworthy that the sphingosine-1-phosphate receptor S1PR3 was upregulated upon SHPS1 silencing. PECA1 silencing led to an increase in levels of AMPN and decreases in levels of SLIT and NTRK-like protein SLIK1, whereas PECA1 was downregulated upon silencing of TRBM or SCRB1. The VEGF-A receptor VEGFR2 was downregulated upon SCRB1 silencing.

To quantitatively assess the interconnectivity of the HDL synapse core candidates, we calculated the fraction of core candidates expected to be affected by the silencing of one candidate. We used the fraction of all HDL synapse core candidates within the total EA.hy926 surfaceome as a reference value. Overall, the fraction of significantly altered HDL synapse core candidates upon stable silencing exceeded the fraction of expected core candidates to be affected (**Suppl. Figure 7F**). This observation supports our hypothesis that the HDL synapse candidates are interconnected. We established an interaction network of all significantly regulated proteins resulting from silencing of PECA1, TRBM, SHPS1, NECT2, and SCRB1 and complemented it with the remaining HDL synapse core candidates and protein-protein interaction evidence from the STRING database (19). This analysis revealed a tightly interconnected cluster of proteins in conjunction with APOA1 (**Figure 3A, Suppl. Table 7**), demonstrating that the endothelial HDL interactome is an interlinked network of receptors that modulate its composition upon intrinsic alterations. To determine the relative importance of these receptors, we first established a STRING database interaction network of the EA.hy926 cell surfaceome (**Suppl. Table 8**). Next, we compared the number of interactions of regulated proteins or core candidates with core candidates to the number of interactions of regulated proteins or core candidates with other EA.hy926 surfaceome proteins (**Suppl. Figure 8**). Sixteen HDL synapse core candidates and six regulated proteins, including SCRB1, VGFR2, PECA1, NECT2, and AMPN, displayed more interactions with core candidates than with other endothelial proteins.

**Figure 3:**
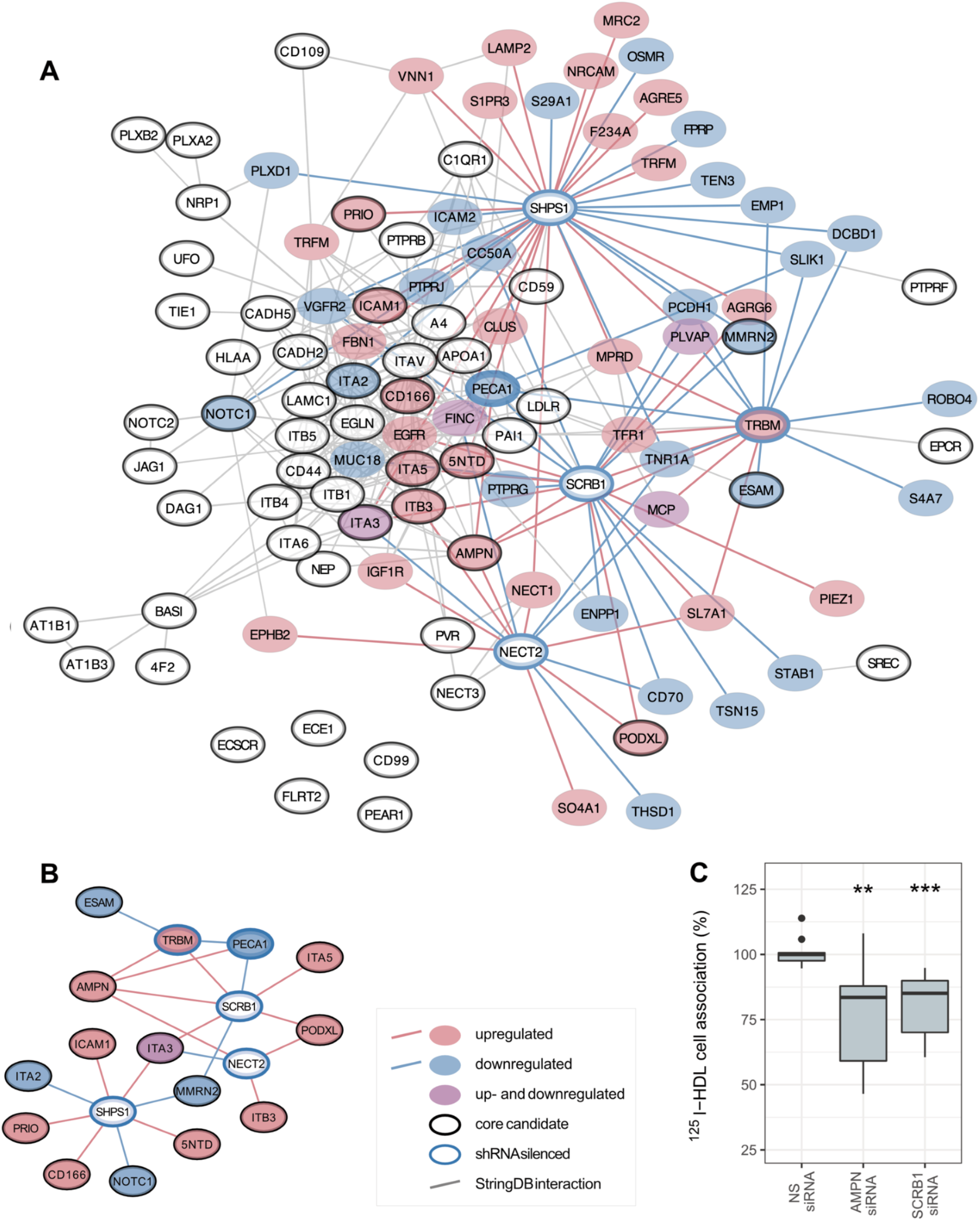
Interconnectivity of endothelial HDL synapse candidates and validation of AMPN as an important member of the interactome. **A)** Interaction network of the endothelial HDL synapse core candidates and proteins that are either up- or downregulated upon stable silencing of PECA1, TRBM, NECT2, SHPS1, or SCRB1. Edges represent either regulated proteins or STRING database confidence >0.7. **B)** Subset of network highlighting endothelial HDL core candidates that are regulated upon stable silencing of PECA1, TRBM, NECT2, SHPS1, or SCRB1 (i.e., FC > 1.5 or < −1.5 and adj. p value < 0.05). **C)** HDL association with HAECs in which either AMPN or SCRB1 were transiently silenced with siRNA. Data are from three independent experiments with quadruplicates or triplicates. Vertical lines indicate the minimum and maximum values, boxes are first and third quartiles, and horizontal lines are medians. Significance was determined with the Wilcoxon rank sum test (** p < 0.01; *** p < 0.001).

### AMPN is a key component of the endothelial HDL synapse

The surface abundance of AMPN, a core candidate of the HDL synapse, was significantly upregulated in four of the five EA.hy926 cell lines deficient in an HDL core candidate, including the line in which SCRB1 was silenced (**Figure 3A and B, Suppl. Figure 7 and 9**). In addition to its broad aminopeptidase activity, AMPN exerts various peptidase-independent receptor functions. For instance, it is a receptor for the human coronavirus 229E (20). As SCRB1 is also known to facilitate the cellular entry of various viruses (21), we hypothesized that AMPN might likewise modulate endothelial HDL uptake. We tested this hypothesis in primary HAECs, in which our HDL-guided LUX-MS experiment also identified AMPN as an HDL-interacting protein (**Suppl. Figure 5A**). First, we assessed the effect of SCRB1 silencing on the AMPN abundance on HAECs. In HAECs in which SCRB1 was transiently silenced with a small interfering RNA (siRNA), the mRNA abundance of AMPN was significantly increased compared to HAECs treated with a control siRNA (**Suppl. Figure 10**). Conversely, the mRNA abundance of SCRB1 was not affected by silencing of AMPN with the same strategy (**Suppl. Figure 10**). siRNA-mediated silencing of AMPN induced moderate surfaceome alterations on HAECs: four proteins were significantly upregulated and three proteins, in addition to AMPN, were downregulated (**Suppl. Figure 11A**). At the protein level, the abundance of SCRB1 was not altered upon silencing of AMPN (**Suppl. Figure 11B**). Silencing of either AMPN or SCRB1 resulted in similar and significant reductions of cellular association with HDL particles (**Figure 3C**). These results indicate that AMPN contributes to HDL uptake into endothelial cells.

### Comparison of endothelial cells and hepatocytes indicates cellspecific HDL synapses

During reverse cholesterol transport, HDL particles engage with several tissues. Hepatocytes take up cholesterol either through SCRB1-mediated selective uptake or HDL holo-particle-mediated uptake for cholesterol recycling and biliary excretion (22). The surfaceomes of endothelial cells and hepatocytes are qualitatively and quantitatively distinct; for example, the surface abundances of SCRB1 differ (13). These differences might influence the HDL interactome. To test this, we investigated if and how the HDL receptor landscapes differ and agree between human hepatocytes, and wild-type EA.hy926 cells using LUX-MS (**Figure 4A, Suppl. Figure 2, Suppl. Table 1**). In HEPG2 cells, the ligand (anti-SCRB1 mouse IgG) and the targeted receptor were both significantly enriched in the anti-SCRB1 antibody-guided LUX-MS experiment, and, in the HDL-based LUX-MS experiment, SCRB1 was significantly enriched. We observed 16 proteins in both the anti-SCRB1-based and the HDL-based LUX-MS-experiments in HEPG2 cells and identified 38 proteins in the proximity of SCRB1 that were not identified by the HDL-based LUX-MS experiment (**Figure 4B**). These results indicate that hepatic SCRB1, unlike endothelial SCRB1, forms additional receptor clusters of different compositions that are not equally targeted by HDL. The binding affinity of HDL is higher to HEPG2 cells than to EA.hy926 cells (**Suppl. Figure 3C-E**), which argues against a more extensive enrichment in EA.hy926 due to affinity and therefore against a technical explanation for the difference in enrichment in the HDL-based LUX-MS experiment on HEPG2 and EA.hy926 cells. Thus, the hepatic HDL synapse might contain a more homogenous receptor community than the endothelial HDL synapse.

**Figure 4:**
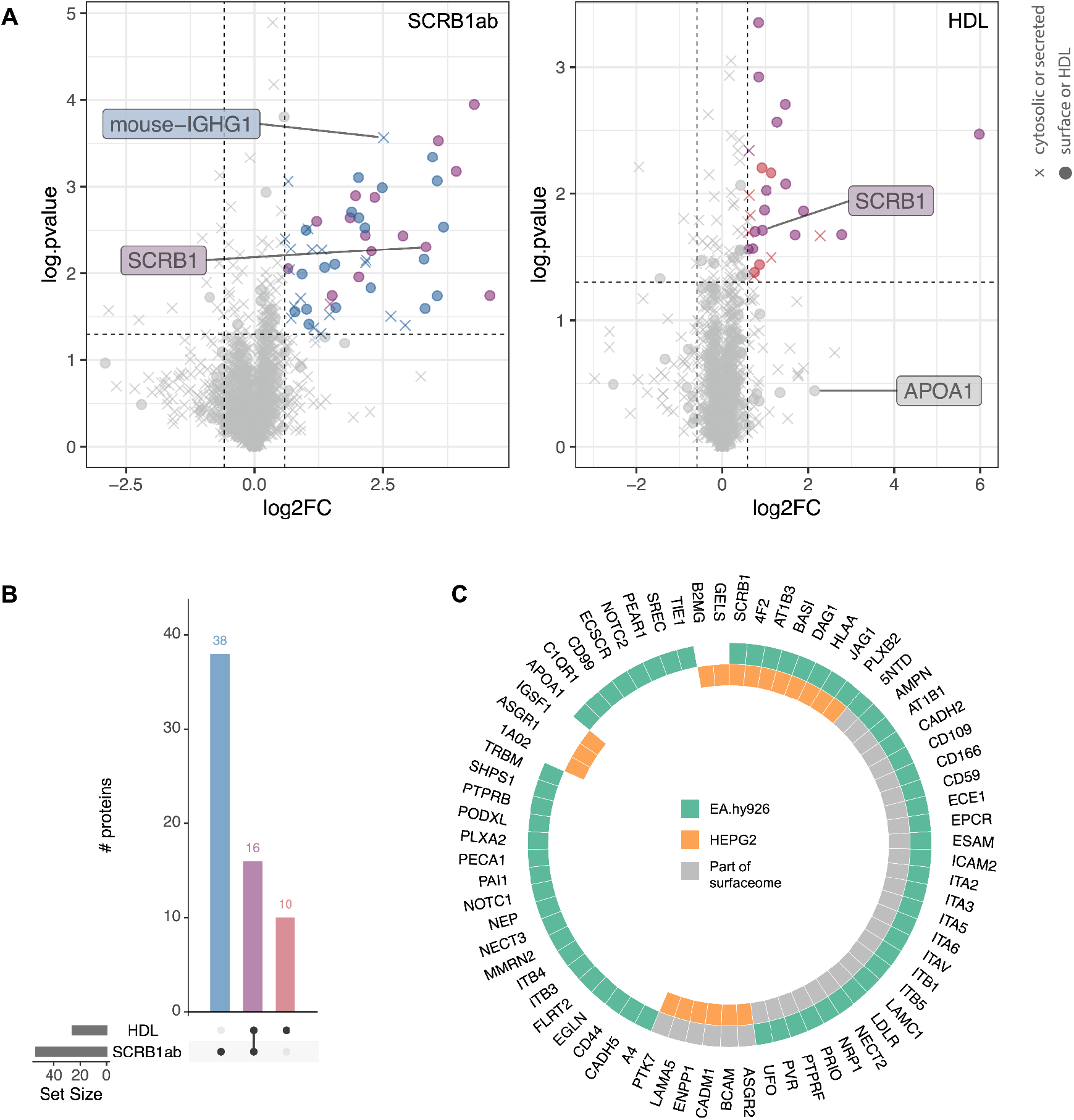
The hepatic HDL synapse and its comparison to the endothelial HDL synapse. **A)** Volcano plots of LUX-MS experiments using an anti-SCRB1 antibody (left) and HDL (right) as a ligand in LUX-MS experiments on HEPG2 cells. Proteins enriched in the anti-SCRB1 antibody-based LUX-MS experiment are in blue. Proteins enriched in the HDL-based LUX-MS experiment are in red. Proteins enriched in both experiments are in purple. **B)** Numbers of proteins detected as enriched in anti-SCRB1 antibody-based and HDL-based LUX-MS experiments and the number of overlapping proteins. **C)** Comparison of the HDL core synapse candidates on EA.hy926 cells (green, outer circle) and the enriched surface/ HDL-associated proteins on HEPG2 cells (orange, inner circle). Grey indicates proteins of the HEPG2 or EA.hy926 surfaceome (13) that were not enriched in the LUX-MS experiment. Significance was determined using MSstats (FC > 1.5 and p value < 0.05). Note: There is no evidence that ECSCR, CD99, or GELS are *N-*glycosylated and therefore the presence on the cell surface cannot be excluded as auto-CSC targets *N-*glycosylated surface proteins.

We next compared the 19 proteins of the hepatic HDL synapse, which are part of Surfy, with the 60 proteins of the endothelial HDL synapse (**Figure 4C**). Eight proteins were shared between the two synapses, including SCRB1. Eleven proteins were exclusively enriched on hepatic cells, and 52 were only enriched on endothelial cells. For instance, the asialoglycoprotein receptors ASGR1 and ASGR2 were exclusively enriched on HEPG2 cells in the HDL-guided LUX-MS experiment. Rare loss-of-function variants of ASGR1 are strongly associated with a reduced risk of coronary heart disease in Icelanders (23). Although present at the cell surface of both EA.hy926 and HEPG2 cells (13), some cell surface proteins only interacted with HDL in the one or the other cell type. AMPN, for instance, is similarly abundant on EA.hy926 and HEPG2 cells (13) but was only enriched in the HDL-based LUX-MS experiments on EA.hy926 cells.

## Discussion

HDL particles are heterogenous complexes of hundreds of different proteins and lipid species (4, 5). Due to this structural diversity and complexity, HDL’s exert many effects on various cell types, which are important for their regular function and survival (6). The cellular interaction partners of HDLs are far from being comprehensively resolved. SCRB1 is the only well-characterized HDL receptor (7). It mediates cholesterol influx and efflux from and to HDL particles, respectively. Also it was found to mediate HDL holoparticle uptake and signal transduction by HDL particles in various cell types (24). SCRB1 is neither necessary nor sufficient for certain HDL functions, however, for example, cholesterol efflux have also been ascribed to ABC transporters A1 and G1 and signal transduction to sphingosine-1-phosphate receptors (10). Redundancy and diversity of HDL interactors likely results from complex cell type-specific networks that interconnect multiple HDL-associated molecules with multiple cell surface proteins. To test the hypothesis that there are many-to-many interactions between HDL particles and cells, we characterized the composition of HDL environment on live cells in a discovery-driven manner using the LUX-MS methodology (11). Our data support the presence of HDL synapses that have cell type-specific spatial organization.

The HDL synapse of wild-type EA.hy926 cells includes SCRB1 and 59 additional proteins. 60% of these core proteins were confirmed by a LUX-MS experiment performed on HAECs, which are primary human aortic endothelial cells. By LUX-MS snapshots of wild-type EA.hy926 cells labeled with heavy amino acids, we validated that APOA1 and six other proteins were derived from the HDL particle. Our previous mass spectrometric studies identified about 300 and about 500 proteins in HDL particles and the EA.hy926 surfaceome, respectively (5, 13); therefore, it was initially surprising that only one of the sixty proteins of the HDL core synapse is exclusively HDL derived, namely APOA1. It is possible that only the minority of the HDL-associated proteins directly interact with cellular counterparts, but it is more likely that current technology is not sensitive enough to capture all interactions. In view of the large range of abundances of HDL-associated proteins (25), it may well be that the number of accessible cell surface proteins versus the number of accessible HDL proteins did not allow us to reproducibly enriche a larger fraction of the HDL proteome at the cell surface. Moreover, it is important to note that the proteome of the HDL particles includes many cellular proteins. Of note, six members of the HDL core synapse, namely CD44, PVR, PAI1, EPCR, ESAM and NRP1, were identified with both heavy and light labels indicating that they were already present in the HDL isolate and also engaging with HDL as part of the EA.hy926 surface. This observation indicates that HDL undergoes cargo exchange at the surface of endothelial cells and, since these proteins are not endothelium-specific, with other types of cells. For example, some of these proteins may have been integrated into the HDL particle during its hepatic or intestinal biogenesis or previous transendothelial passages.

We perturbed the cell surface abundances of four HDL synapse proteins and SCRB1 on EA.hy926 cells by stable silencing. The silencing of single proteins led to the quantitative reorganization of HDL interacting cell surface proteins. HDL synapse core proteins identified by the HDL-based LUX-MS experiments and regulated proteins appear to form a tightly connected interaction network. We hypothesize that members of the HDL synapse play three distinct roles. Proteins of the first group directly interact with HDL particles, as previously shown for SCRB1 (7) and as suggested for the CD166 antigen (which is encoded by *ALCAM)* (26). Both of these receptors were enriched in the HDL-guided LUX-MS snapshots of EA.hy926 cells and HAECs. The second group might maintain the overall receptome integrity. As an example of a protein that functions in this manner, SHPS1 was suggested to serve as a scaffold for multi-protein complex assembly to modulate adhesion-regulated signals in macrophages (27). The HDL surfaceome underwent major quantitative reorganization upon SHPS1 silencing, suggesting that SHPS1 might serve as a general scaffold within the HDL synapse. Proteins of the third group interact transiently with HDL but may nevertheless be functionally relevant. For instance, EPCR, enriched in the HDL-guided LUX-MS snapshot of both EA.hy926 and HAEC, and also carried by HDL, is known to regulate coagulation with TRBM, an HDL-interacting protein in both endothelial cell types, through activation of protein C (28). Protein C, in turn, utilizes HDL as a cofactor to mediate anticoagulation and endothelial barrier protection (29). EPCR was suggested to exert cytoprotective functions through PAR1, angiopoietin-1 receptor TIE2, and S1PR1 (30). Most likely due to the sparsity of accessible reaction sites for LUX-MS, we did not identify PAR1 nor S1PR1 in our interactomes. S1PR1 was found to transiently interact with SCRB1 through a mechanism mediated by HDL binding, to trigger calcium flux and S1PR1 internalization (31). TIE2 was enriched in the HDL-based LUX-MS experiments on EA.hy926 cells that overexpress SCRB1 and in the LUX-MS snapshot of HAEC. On the other hand, TIE1 is part of the EA.hy926 HDL synapse core candidate list. TIE1 is thought to modulate TIE2 activity and thereby angiogenesis (32). Silencing of NRP1, a cofactor of VGFR2, was previously shown to reduce lipoprotein uptake in clear-cell renal cell carcinomas cells, which accumulate lipids in a VEGF- and SCRB1-dependent manner (33). VGFR2 was not identified in anti-SCRB1 antibody or HDL-based LUX-MS experiments. However, on both endothelial cell types, we identified NRP1 and multimerin MMRN2, which was suggested to interfere with VGFR2 activation by sequestering VEGF-A (34). Moreover, we identified PTPRB, which was shown to dephosphorylate VGFR2 (35). Furthermore, VEGFR2 was downregulated when SCRB1 expression was silenced.

AMPN was identified as a core protein of the HDL synapse of both EA.hy926 cells and HAECs. AMPN was not detected in the HDL-guided LUX-MS experiment on HEPG2 cells, although it is present on these cells (13). AMPN was upregulated upon silencing of SCRB1, PECA1, THBR, and NECT2 in EA.hy926 cells. The abundance of AMPN was also increased on the HAEC surface upon SCRB1 silencing. Therefore, AMPN might have redundant functions with these proteins. AMPN is a multifunctional receptor with broad peptidase activity and other peptidase-independent functions. For instance, it was shown to regulate monocyte adhesion to endothelial cells (36). We showed that loss of AMPN compromises HDL uptake into HAECs. AMPN levels are significantly reduced in HDL particles derived from patients with type 2 diabetes compared to HDL derived from the healthy cohort (5). This is additional evidence that HDL might interact with AMPN and that this receptor might play an important HDL-related role.

Several endothelial HDL synapse core proteins were associated with gene ontology terms related to virus-host interactions. SCRB1 participates in hepatitis C virus cell entry (37) and facilitates cell entry of SARS-CoV-2 bound to HDL (38). NRP1, another core protein of the endothelial HDL synapse, also contributes to SARS-CoV-2 entry into host cells (39, 40). PVR is a poliovirus receptor (41), and NECT2 was reported to be critical for herpes simplex virus entry (15). Finally, AMPN is a receptor for the human coronavirus 229E (20). These data suggest that virus particles hitchhike HDL endocytosis pathways, possibly because of their similar molecular interaction faces, which are composed of proteins and lipids.

In HDL-guided LUX-MS experiments, not all proteins in the EA.hy926 surfaceome were detected in the HEPG2 surfaceome and vice versa. This indicates that different cell types might form distinct specialized receptor clusters, targeted explicitly by a subset of HDL particles. SCRB1 was captured by the HDL-based LUX-MS experiment in both EA.hy926 and HEPG2 cells. However, we observed two different enrichment patterns when we compared HDL-guided and anti-SCRB1 antibody-guided LUX-MS snapshots on these cell types. Interestingly, the endothelial SCRB1 environment almost entirely encompassed the HDL interaction space, whereas the hepatic SCRB1 environment was only partly targeted by HDL. On hepatic cells, SCRB1 might form different nanoscale organizations that conduct HDL-dependent and HDL-independent functions. Hepatic SCRB1 is mainly involved in selective lipid uptake from HDL, whereas endothelial SCRB1 facilitates holoparticle uptake of both HDL and LDL and transduces signals that, for example, activate eNOS (8, 22). Thus, the different modes of action in response to binding of HDL to SCRB1 on hepatic or endothelial cell surfaces might require different interaction partners. Furthermore, endothelial cells have a predominant glycocalyx layer (42) that might lead to additional enrichment of extracellular matrix proteins. Indeed, 20 endothelial HDL synapse members were associated with the gene ontology term extracellular matrix organization.

In conclusion, the presented data suggest that cell type-specific spatial receptor organizations are involved in HDL signaling and that these organizations can be characterized using our HDL-guided LUX-MS strategy. Our findings support the hypothesis that the HDL synapse is an interlinked organization of receptors that modulate HDL binding, uptake, and signaling. In the future, our data will serve as a resource for deciphering complex interactions between HDL particles and cells and the underlying functions of these interactions in targeted molecular biology approaches. Eventually, understanding the HDL structure-function mosaic will pave the way for new treatment and prevention strategies for a wide range of pathologies.

## Abbreviations

(auto-CSC): Automated Cell Surface Capturing
(Human endothelial somatic hybrid cells): EA.hy926
(human hepatocellular carcinoma): HEPG2
(HDL): High-density lipoproteins
(HAEC): Human aortic endothelial cells
(SOG): Singlet oxygen generators
(siRNA): Small interfering RNA

## List of human genes and protein names discussed in the paper

ALCAM: CD166 antigen (CD166)
ANPEP: Aminopeptidase N (AMPN/ CD13)
APOA1: Apolipoprotein A (APOA1)
ASGR1: Asialoglycoprotein receptor 1 (ASGR1)
ASGR2: Asialoglycoprotein receptor 1 (ASGR2)
ATP1B1: Sodium/potassium-transporting ATPase subunit beta-1 (AT1B1)
ATP1B3: Sodium/potassium-transporting ATPase subunit beta-1 (AT1B3)
AXL: Tyrosine-protein kinase receptor UFO (UFO)
DAG1: Dystroglycan (DAG1)
ESAM: endothelial cell-selective adhesion molecule (ESAM)
F2R: Proteinase-activated receptor 1 (PAR1)
ICAM1: Intercellular adhesion molecule 1 (ICAM1)
KDR: Vascular endothelial growth factor receptor 2 (VGFR2)
LDLR: Low-density lipoprotein receptor (LDLR)
MMRN2: Multimerin-2 (MMRN2)
NECTIN2: Nectin-2 (NECT2/ PVRL2)
NRP1: Neuropilin-1 (NRP1)
PECAM1: Platelet endothelial cell adhesion molecule (PECA1)
PROCR: Endothelial protein C receptor (EPCR)
PTPRB: Receptor-type tyrosine-protein phosphatase beta (PTPRB)
PTPRF: Receptor-type tyrosine-protein phosphatase F (PTPRF)
PVR: Poliovirus receptor (PVR)
S1PR1: Sphingosine-1-phosphate receptor 1 (S1PR1)
S1PR3: Sphingosine 1-phosphate receptor 3 (S1PR3)
SCARB1: Scavenger receptor B1 (SCRB1)
SCARF1: Scavenger receptor class F member 1 (SREC)
SERPINE1: plasminogen activator inhibitor 1 (PAI1)
SIRPA: Tyrosine-protein phosphatase non-receptor type substrate 1 (SHPS1)
SLC3A2: 4F2 cell-surface antigen heavy chain (4F2)
SLITRK1: SLIT and NTRK-like protein 1 (SLIK1)
TEK: Angiopoietin-1 receptor (TIE2)
THBD: Thrombomodulin (TRBM)
TIE1: Tyrosine-protein kinase receptor Tie-1 (TIE1)
VEGFA: vascular endothelial growth factor A (VEGF-A)

## Contributions

If not mentioned otherwise, experiments and data analysis were performed by KF. HDL-binding and uptake assays were performed by LR. AP created the EA.hy926 V1 overexpressing cells. AvE, LR, and SG provided expertise and critical feedback at all stages of the project. KF and BW conceived the project and designed experiments. KF wrote the first and subsequent versions of the paper. All authors critically read and revised the manuscript.

## Acknowledgements

We thank Silvija Radosavljevic and Eveline Schlumpf for their help with HDL isolation. We acknowledge the BW research group for support at all stages of the project. We thank J. R. Wyatt for text editing. SG was supported by funding through the Personalized Health and Related Technologies strategic focus area of ETH. This work was supported by ETH grant 30 17-1 (to BW), the Swiss National Science Foundation (grant 31003A_160259 to BW), a SystemsX.ch special opportunity grant (to BW), a SystemsX.ch MRD grant (2014/267 to AvE and BW), and the Swiss National Science Foundation (310030-185109 to AvE).

## Materials and methods

### General information

All standard chemicals, if not indicated otherwise, were purchased from MERCK.

### Mammalian cell culture

EA.hy926 cells (ATCC CRL-2922^™^) and HEPG2 cells (ATCC^®^ HB-8065^™^) were grown in Dulbecco’s Modified Eagle’s Medium (DMEM, Gibco) containing 10% fetal bovine serum (FBS, Amimed) and 1% penicillin-streptomycin (Life Technologies Europe B.V.) supplemented with high glucose (4.5 mg/l) and GlutaMax. For isotope labeling experiments, EA.hy926 cells were grown in SILAC DMEM high glucose with stable glutamine and without arginine and lysine (Pan Biotech), supplemented with 10% dialyzed FBS (Pan Biotech) and 1% penicillin-streptomycin (Life Technologies Europe B.V.), 200 mg/L L-proline, 50 mg/L L-arginine[10], and 50 mg/L L-lysine[8] (Silantes) for 20 days. The incorporation rate of heavy amino acids was determined to be 99% by MS analysis of the lysate, whereas the incorporation rate for surface proteins was 95%. HAECs (Cell Applications Inc (304-05a)) were cultured in Human Endothelial Cell Growth Medium, All-in-one ready-to-use (Cell Applications Inc, 211A-500) without antibiotics.

### HDL isolation and processing

HDL (1.063<d<1.21 g/mL) was isolated from fresh human normolipidemic plasma of blood donors by sequential ultracentrifugation as described previously (43). For LUX-MS experiments SOGs were coupled to HDL particles. HDL particles were diluted in coupling buffer (95% PBS and 5% 0.2 M sodium bicarbonate, pH 8.3) and NHS-functionalized thiorhodamine (Thio12, cat: AD-Thio12-41, ATTO-TEC GmbH) was added to an approximate final molecular weight ratio of 1:25 HDL to SOG. The mixture was incubated for 1 h at room temperature with shaking at 300 rpm in the dark. Subsequently, the buffer was exchanged to PBS using 7k MWCO Zeba Spin Desalting Columns (ThermoFisher Scientific). For the flow cytometry experiments, HDL particles were labeled with Atto 488 NHS (ATTO-TEC GmbH). In brief, 1 mg HDL preparation was diluted in PBS, pH 8 and mixed with 50 μg Atto 488 NHS. After an incubation of 1 h, the HDL-Atto 488 construct was purified using a PD-10 column.

### HDL-guided and anti-SCRB1 antibody-guided LUX-MS

The LUX-MS experiments were performed as previously described (11). Per replicate, one 10-cm petri dish containing approximately 10 million cells confluent EA.hy926 cells, HEPG2 cells, or HAECs was used. For all LUX-MS conditions, triplicates were generated. Cells were precooled for 20 min at 4 °C and then incubated with HDL-SOG for 40 min at 4 °C with 50 μg HDL per ml PBS or with anti-SCRB1 antibody-SOG (BioLegend, 363201; antibody concentration 1:100; antibody-SOG ratio 1:5) for 15 min at 4 °C. The control samples were treated with the same amount of HDL or mouse IgG Isotype control antibody (Invitrogen, 10400C) not coupled to SOG. After incubation, cells were washed three times with ice-cold PBS to remove unbound HDL-SOG, HDL, anti-SCRB1 antibody, or the isotype control. Subsequently, pre-cooled photo-oxidation buffer (5 mM byocytin hydrazide (Pitsch Nucleic Acids) in heavy water-based PBS (deuterium oxide (ARMAR AG, 014100.2065)), pH 7.5) was added. The plates were illuminated with 590 nm Precision LED spotlights controlled by a BioLED Light Control Module (Mightex Systems). After 15 min of illumination, the photo-oxidation buffer was removed and pre-cooled labeling buffer (5 mM byocytin hydrazide and 50 mM aminomethyl imidazole dihydrochloride in PBS, pH 6) was added. Cells were incubated for 1 h at 4 °C and subsequently washed three times with ice-cold PBS followed by scraping and an additional washing step by centrifugation for cell pelleting. Cells were then lysed in lysis buffer (100 mM Tris, 1% sodium deoxycholate, 10 mM TCEP, 15 mM 2-chloroacetamide) by repetitive sonication using a VialTweeter (Hielscher Ultrasonics) and heated at 99 °C for 5 min. Protein content was determined using a NanoDrop 2000 instrument (ThermoFisher Scientific), and samples were normalized accordingly before capture of biotinylated proteins on Pierce^™^ Streptavidin UltraLink^™^ Resin (80 μl; ThermoFisher Scientific), washing, and digestion with lysyl endopeptidase (Lys-C, Wako) and sequencing-grade trypsin (Promega). The capture and digestion steps were performed with a Versette liquid handling robot (ThermoFisher Scientific). Sample purification was performed over C18 resin (100 μg capacity; The Nest Group) using 5% to 80% acetonitrile, 0.1% formic acid (FA; Chemie Brunschwig).

### auto-CSC of plasma membrane proteins

Cell surface capturing of EA.hy926 cells in which TRBM, PECA1, SCRB1, NECT2, or SHPS1 were stably silenced and of cells that express the control shRNA was performed as previously described (12). In brief, around 20 million cells were used per sample. For each condition, triplicates were generated. Cells were oxidized with 3 mM sodium periodate in PBS with 0.1% FBS at pH 6.5 for 15 min at 4 °C in the dark and labeled with 4 mM biocytin hydrazide and 50 mM aminomethyl imidazole dihydrochloride in PBS at pH 7.4 for 1 h at 4 °C. After every step, cells were washed with ice-cold PBS. Cells were harvested by scraping, lysed in lysis buffer by repetitive sonication using a VialTweeter (Hielscher Ultrasonics), and heated at 99 °C for 5 min. Proteins were digested and biotinylated peptides were captured (80 μl Pierce^™^ Streptavidin UltraLink^™^ Resin, ThermoFisher Scientific), resin was washed, and peptides were released by PNGaseF (BioConcept AG) on the Versette liquid handling robot (ThermoFisher Scientific). The unbound samples were used for the assessment of silencing efficiency. Sample purification was performed over C18 resin (100 μg capacity; The Nest Group) using 5% to 80% acetonitrile, 0.1% FA (Chemie Brunschwig).

### LC-MS/MS

Peptide samples were reconstituted in 3% acetonitrile with 0.1% FA in HPLC-grade water. The samples (1 μg) were loaded onto an EASY-nano-HPLC system (EASY-nLC 1000 or 1200, ThermoFisher Scientific) mounted with a reverse-phase HPLC column (75 μm ID) in-house packed with 15 cm stationary phase (Reprosil-Pur C18-AQ 1.9 μm, 200 Å, Dr. Maisch). The HPLC was associated with either a Q Exactive Plus Mass Spectrometer (ThermoFisher Scientific) or a Q Exactive HF Mass Spectrometer (ThermoFisher Scientific) and a nano-electrospray ion source (ThermoFisher Scientific).

For LUX-MS-derived peptides measured on the Q Exactive Plus Mass Spectrometer, peptides were loaded onto the column with buffer A (99.9% HPLC grade water, 0.1% FA) and eluted with a 60-min gradient of 5-30% buffer B (99.9% acetonitrile, 0.1% FA) followed by a 5-min gradient from 30-50% buffer B, and two washes with 95% buffer B. For LUX-MS-derived peptides and whole-cell lysates measured on the Q Exactive HF Mass Spectrometer, peptides were loaded onto the column with buffer A (99.9% HPLC grade water, 0.1% FA) and eluted with a 60-min gradient of 6-36% buffer B (80% acetonitrile, 19.9 % HPLC grade water, 0.1% FA) followed by a 5-min gradient from 36-60% buffer B, and two washes with 100% buffer B. The auto-CSC-derived peptides were measured on a Q Exactive HF Mass Spectrometer. Peptides were loaded onto the column with buffer A (99.9% HPLC grade water, 0.1% FA) and eluted with a 45-min gradient of 3-44% buffer B (80% acetonitrile, 19.9 % HPLC grade water, 0.1% FA) followed by a 4-min gradient from 44-60% buffer B, and two washes with 98% buffer B.

Mass spectra were acquired in data-dependent acquisition (DDA) mode (Top15). MS1 spectra were acquired at 60,000 resolution in the range of 375 to 1,700 m/z, for the Q Exactive HF Mass Spectrometer or 70,000 resolution in the range of 300 to 1,700 m/z for the Q Exactive Plus Mass Spectrometer. The automatic gain control target value was 3×10^6^, and the maximum injection time was set to either 110 or 120 ms. MS/MS was acquired with a resolution of 15,000 or 35,000. Precursor ions were fragmented using HCD with an isolation width of 1.2 or 1.4 m/z and an automatic gain control target value of 10^5^ or 10^6^. Target ions already selected for fragmentation were dynamically excluded for 15 s or 30 s.

For HDL proteotyping, peptides were measured on the Q Exactive HF Mass Spectrometer. Peptides were loaded onto the column with buffer A (99.9% HPLC grade water, 0.1% FA) and eluted with a 60-min gradient of 5-41% buffer B (80% acetonitrile, 19.9 % HPLC grade water, 0.1% FA) followed by a 5-min gradient from 41-52% buffer B, and two washes with 98% buffer B. Mass spectra of HDL protein samples and cell lysates, containing retention time reference peptides (iRT, Biognosys), were acquired in data-independent acquisition (DIA) mode. After a survey scan acquired in the Orbitrap with 120,000 resolution, 60 ms max injection time, and an automatic gain control target of 3×10^6^, 24 DIA segments followed with a stepped collision energy of 25%, 27%, and 30% acquired in the Orbitrap with 30,000 resolution, injection time set to auto, and the automatic gain control target set to 3×10^6^. The mass range was set to 350 - 1650 m/z.

### Mass spectrometry data analysis

Files were converted into mzML using MSconvert for the DDA-based RAW data. Using COMET (v27.0), fragment ion spectra were searched against UniprotKB (Swiss-Prot, Homo sapiens from March 2019) containing common MS contaminants (without bovine apolipoproteins) and standards. For SCRB1 the sequences of all variants were added due to a discrepancy between UniProt and the common naming convention for variant 1 (21). The precursor mass tolerance was set to 20 ppm. Search parameters were fully-tryptic and semi-tryptic for auto-CSC samples with carbamidomethylation as a fixed modification for cysteines. Oxidation of methionine was set as a variable modification. For auto-CSC samples, deamidation of asparagine was also set as a variable modification. The PeptideProphet probability scoring was calculated using the Trans-Proteomic Pipeline (v4.6.2). Peptides with an error rate of ≤ 1% were selected for quantification. Peptide identifications were further filtered for the presence of the consensus NXS/T sequence with simultaneous deamidation (+0.98 Da) at asparagines for auto-CSC samples. For label-free MS1-based quantification, non-conflicting peptide intensities were extracted in Progenesis QI (Nonlinear Dynamics). DIA measurements of the HDL samples and the lysates were analyzed with Spectronaut Pulsar version 14 (Biognosys) using default settings. The previously established HDL spectral library was used for HDL proteotyping (5). For the lysates, a library was created using Proteome Discoverer v2.4 (ThermoFisher Scientific). The spectra were searched against UniprotKB (Swiss-Prot, Homo sapiens from March 2019) containing common MS contaminants (without bovine apolipoproteins) and standards using the Sequest HT search engine within Proteome Discoverer v2.4. The names reported are the UniProt main names without the “_HUMAN” add-on. All full names, frequently used alternative names, accessions, and gene names of the proteins that are part of a main figure can be found in **Suppl. Table 9**. Mass spectrometry raw data were deposited on the MassIVE data repository under the identifier MSV000090734.

### Statistical data evaluation and visualization

For the calculation of the protein abundance changes and statistical data evaluation for the auto-CSC and the LUX-MS experiments, Progenesis results, or Spectronaut results in case of the cell lysates, were processed with MSstats (v3.8.6.) (44) in the R computing environment (v4.0.2.) with default settings. In brief, protein abundance differences were determined with a linear mixed effect model and tested for statistical significance using a two-sided t-test. For **Figure 2**, minimal feature settings were set to 1 per protein per condition (instead of 2), due to the cumulative evidence from four experiments, but excluded if at least two features were not detected across the four experiments. Cell surface localization was assigned using the *in silico* human surfaceome (14) and the previously established auto-CSC snapshots of EA.hy926 cells, HAECs, and HEPG2 cells (13). HDL annotation was assigned using the previously established inventory of HDL proteins (5) as well as the Davidson HDL watch list (http://homepages.uc.edu/~davidswm/HDLproteome.html, March 2021). For the HDL proteotypes, the data matrix was exported from Spectronaut Pulsar v14 (Biognosys). The data were processed in R (v4.0.2.) and visualized with the ggplot2 and the UpSetR packages. The networks in **Figure 3** were created with Cytoscape V3.8.2 (45) using STRING database data (https://string-db.org/). The gene ontology analysis was performed with the Cytoscape plugin ClueGo using the standard settings (46, 47). Illustrations in **Figure 1 A** were created with BioRender.com and the Servier Medical Art collection (https://smart.servier.com).

### Flow cytometry

To assess HDL binding by flow cytometry, 50 ug/ml HDL-Atto 488 was added to the EA.hy926 cells, EA.hy926 cells that overexpress SCRB1 V1, EA.hy926 cells with silenced SCRB1, or HEPG2 cells and incubated for 40 min at 4 °C. For cytometric cell surface SCRB1 abundance estimation, cells were incubated with the anti-SCRB1 antibody (BioLegend, 363201) for 15 min at 4 °C. To reduce the amount of antibody used, cells were scraped before primary antibody incubation. Subsequently, the cell-bound anti-SCRB1 antibody was labeled with Goat Anti-Mouse IgG DyLight (ThermoFisher Scientific, 35503) for 15 min at 4 °C. For cytometric LUX-MS cell surface biotinylation assessment, biotinylated cells were labeled with NeutrAvidin Dylight 488 (ThermoFisher Scientific, 22832). After every incubation step, the cells were washed with ice-cold PBS. The viability of cells was determined using propidium iodide staining, samples were measured on a BD Accuri C6 Cytometer (BD Biosciences), and the acquired data were analyzed and visualized using the FlowJo software (v.10.07).

### Establishment of stable and transient gene silencing or overexpression

For gene silencing in EAhy926 cells, a lentiviral shuttle plasmid for shRNA targeting human *SCARB1* (TRCN0000056966, pLKO.1), human *THBD* (TRCN0000053923, pLKO.1), human *NECTIN2* (TRCN0000063075, pLKO.1), human *PECAM1* (TRCN0000057801, pLKO.1) or human *SIRPA* (TRCN0000002684, pLKO.1) was used. The control was Eahy926 cells transfected with a control plasmid (a gift from David Sabatini (Addgene plasmid #1864; http://n2t.net/addgene:1864; RRID:Addgene_1864)) (48). The transduction was enhanced with Polybrene (8 μg/ml), and cells were selected with puromycin for three passages before the experiments. The extent of silencing was assessed with RT-qPCR (**Suppl. Figure 6A**), flow cytometry (**Suppl. Figure 1C and D**), auto-CSC (**Suppl. Figure 6B**), and in the cell lysates (**Suppl. Figure 6C**). Lentiviruses were packaged according to the following protocol: Shuttle plasmid (8 μg), psPAX2 packaging vector (2 μg, a gift from Didier Trono (Addgene plasmid #12260; http://n2t.net/addgene:12260; RRID: Addgene_12260)), and pMD2G envelope plasmid (4 μg, a gift from Didier Trono (Addgene plasmid #12259; http://n2t.net/addgene:12259; RRID: Addgene_12259)) were transfected into HEK-293 T cells (ThermoFisher Scientific) with 1:3 DNA: polyethylenimine. The medium was exchanged after 12 h with DMEM containing 10% FBS and 1% penicillin-streptomycin. After 48 h, the supernatant was collected, filtered through a 0.2-μm filter (Sarstedt AG), and aliquoted.

For silencing of AMPN, HAECs were transfected with siRNA targeted to *ANPEP* or *SCARB1* or with a non-silencing control (Dharmacon, SMARTpool) at a final concentration of 5 nM using Lipofectamine RNA iMAX transfection reagent (Invitrogen, 13778150) in an antibiotic-free medium. The extent of silencing was assessed with RT-qPCR (**Suppl. Figure 10**), auto-CSC (**Suppl. Figure 11A**), and in cell lysates (**Suppl. Figure 11B**). All experiments were performed 72 h post-transfection. The cDNA encoding SCRB1 variant 1 (Uniprot accession: Q8WTV0-2), cloned into expression vector pReceiver M02, was purchased from Genecopoeia. Using Lipofectamine 2000 (ThermoFisher Scientific), the construct or pReceiver M02 without insert were transfected into EA.hy926 cells. The cells were selected for three passages in DMEM supplemented with 10% FCS and 500 μg/mL G418 sulfate (ThermoFisher Scientific, 10131027). The extent of overexpression was assessed with flow cytometry (**Suppl. Figure 1A and B**).

### Quantitative real-time PCR for validation of silencing efficiency

Total RNA was isolated using the RNeasy Mini Kit (QIAGEN, 74104) according to the manufacturer’s instructions. Genomic DNA was removed by digestion using DNase (Roche) in the presence of Ribolock RNase inhibitor (ThermoFisher Scientific). Reverse transcription was performed using 200 U/μL M-MLV RT (Invitrogen) following the standard protocol as described by the manufacturer. Quantitative PCR was performed with Lightcycler FastStart DNA Master SYBR Green I (Roche) using gene-specific primers listed in **Suppl. Table 10**. CP values were fit to the standard curve and normalized to *GAPDH*.

### HDL association experiments

Quantification of cell association of radiolabeled HDL particles to HAECs was performed as previously described (49). The experiments were performed in DMEM (Sigma) containing 25 mM HEPES and 0.2% BSA. Cells were incubated with 10 μg/mL of ^125^I-HDL without or with 40-fold excess of unlabeled HDL for 1 h at 37 °C. The specific cellular association was calculated by subtracting the values obtained in the presence of excess unlabeled HDL from those obtained in the absence of unlabeled HDL.

