## Supplementary Figures for "Mapping the dynamic cell surface interactome of high-density lipoprotein reveals Aminopeptidase N as modulator of its endothelial uptake"

### Supplementary information

#### Supplementary tables:

**Suppl. Table 1:** Significant testing results (MSstats export) of all LUX-MS snapshots. This table includes light to heavy labeled peptide intensity ratios if applicable.

**Suppl. Table 2:** Protein abundance (Spectronaut output) of the different HDLs used as ligands for four independent LUX-MS snapshots on EA.hy926.

**Suppl. Table 3:** Peptide quantification data table of the two LUX-MS snapshots on heavy labeled EA.hy926 cells (MSstats output).

**Suppl. Table 4:** Summary table of the gene ontology analysis of the EA.hy926 HDL synapse core candidates.

**Suppl. Table 5:** Significant testing results (MSstats export) of the auto-CSC experiments on EA.hy926 cells with shRNA-silenced genes or on HAEC with siRNA-silenced AMPN.

**Suppl. Table 6:** Protein abundance differences from lysates of the cell lines with silenced genes compared to the NS control cell line.

**Suppl. Table 7:** STRING DB interactions (confidence > 0.7) as well as connection evidence (FC > 1.5, adj. p value < 0.05) from the auto-CSC with shRNA silenced genes.

**Suppl. Table 8:** STRING DB interactions (confidence > 0.7) of the whole EA.hy926 surfaceome established with auto-CSC.

**Suppl. Table 9:** All full names, frequently used alternative names, accessions, and gene names of the proteins that are part of a main figure or supplementary figure 4.

**Suppl. Table 10:** List of primers used for RT-qPCR.

#### Supplementary figures:

**Suppl. Figure 1:** SCRB1 cell surface abundance analysis on SCRB1 overexpressing or silenced EA.hy926 by flow cytometry.

**Suppl. Figure 2:** Overview of all enriched proteins in anti-SCRB1 antibody and HDL-based LUX-MS snapshots across all different cell lines.

**Suppl. Figure 3:** Flow cytometry-based analysis of HDL binding rates on different cell types.

**Suppl. Figure 4:** Protein abundance ranking plot of the different HDLs used as ligands for four independent LUX-MS snapshots on EA.hy926.

**Suppl. Figure 5:** Validation of the endothelial HDL synapse on HAEC.

**Suppl. Figure 6:** Abundance validation of shRNA silenced genes on EA.hy926.

**Suppl. Figure 7:** Quantitative surfaceome analysis of EA.hy926 cells upon shRNA mediated silencing of

**Suppl. Figure 8:** Comparison of the number of interactions of regulated or core candidates with core candidates against the number of interactions with other EA.hy926 surfaceome proteins.

**Suppl. Figure 9:** AMPN mRNA levels upon stable SCRB1 silencing in EA.hy926.

**Suppl. Figure 10:** Relative mRNA abundance of AMPN and SCRB1 upon siRNA silencing of AMPN and SCRB1 in HAEC.

**Suppl. Figure 11:** Abundance validation of siRNA silenced AMPN on HAEC and surfaceome dynamics.

A

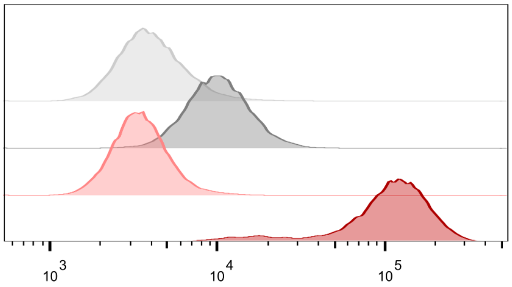


| | Workspace Annotation | Median : FL1-H |
|---|---|---|
| | Vector control + mIg-DL488 | 3912 |
| | Vector control + antiSCRB1ab + mIG-DL488 | 10025 |
| | SCRB1 overexpressing + mIgG-DL488 | 3422 |
| | SCRB1 overexpressing + antiSCRB1ab + mIgG-DL488 | 52995 |

C

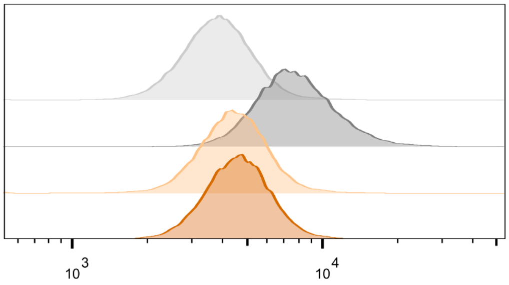


| | Workspace Annotation | Median : FL1-H |
|---|---|---|
| | NS shRNA + mIgG-DL488 | 3799 |
| | NS shRNA + antiSCRB1ab + mIgG-DL488 | 7646 |
| | SCRB1 shRNA + mIgG-DL488 | 4402 |
| | SCRB1 shRNA + antiSCRB1ab + mIgG-DL488 | 4589 |

B

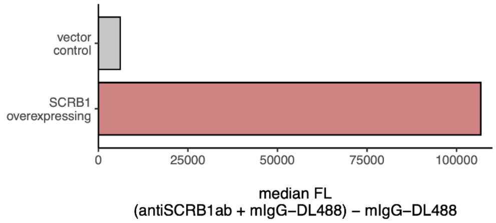


D

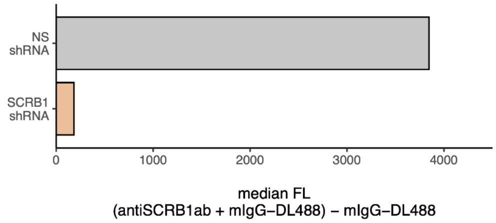


**Suppl. Figure 1:** SCRB1 cell surface abundance analysis on EA.hy926 cells that overexpress SCRB1 or in which SCRB1 expression was silenced. **A)** Flow cytometry analyses of SCRB1 abundance on EA.hy926 cells transfected with vector for expression of control shRNA compared to EA.hy926 cells in which SCRB1 was silenced by expression of shRNA. **B)** Median fluorescence signals of anti-SCRB1 antibody detected with labeled secondary antibody subtracted from control incubated with secondary antibody only. **C)** Flow cytometry analyses of SCRB1 abundance on vector control-transfected EA.hy926 cells and SCRB1-overexpressing EA.hy926 cells. **D)** Median fluorescence signals of the anti-SCRB1 antibody detected with labeled secondary antibody subtracted from control incubated with secondary antibody only.

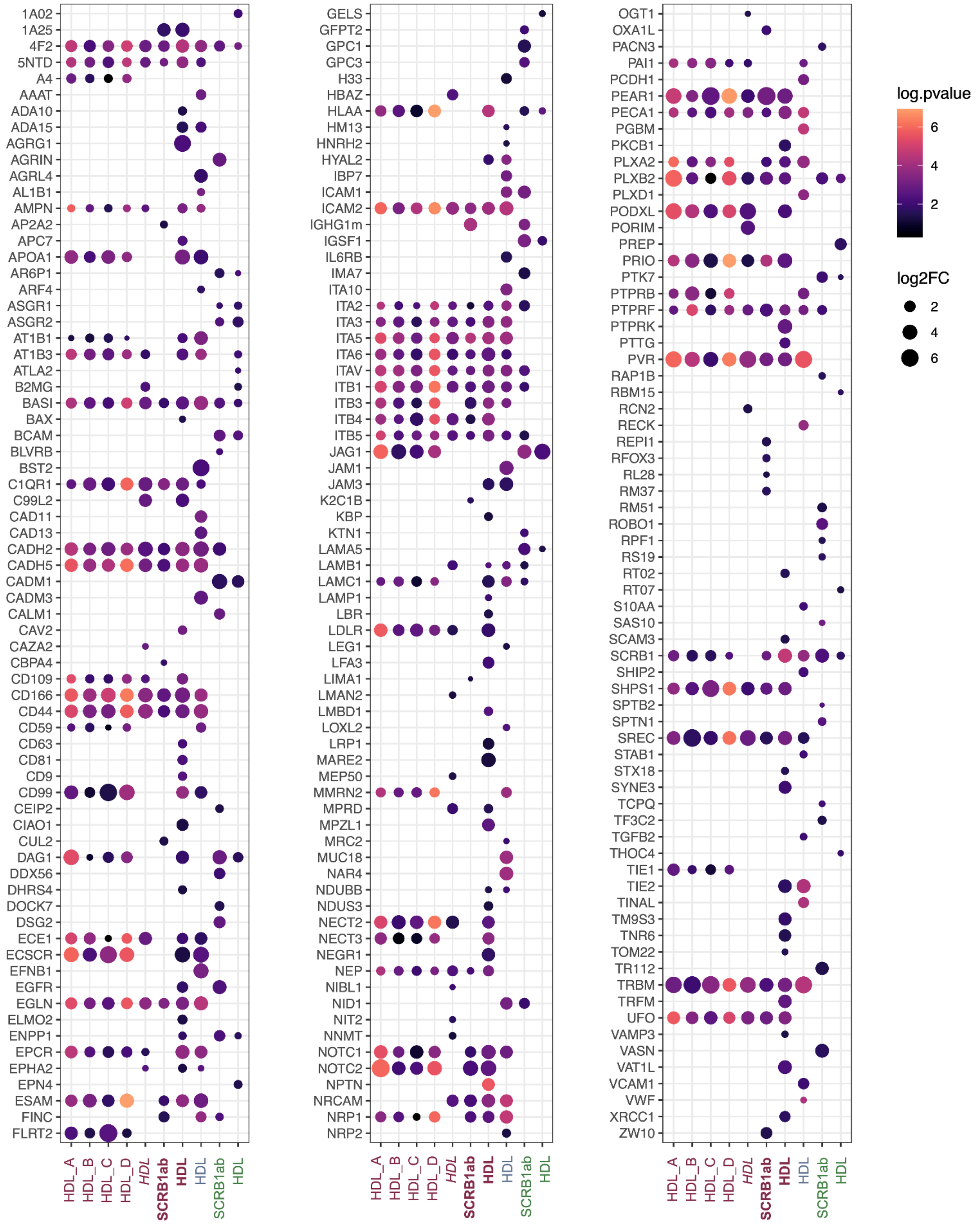


**Suppl. Figure 2:** Overview of all enriched proteins in anti-SCRB1 antibody-based and HDL-based LUX-MS snapshots in indicated cell lines. HDL_A, HDL_B, HDL_C, and HDL_D display the 60 HDL synapse core candidates. For the other snapshots all significant proteins are included (i.e., FC > 1.5 and p value < 0.05). Dot sizes correspond to FC and dot colors indicate p values. X-axis labels: Red: EA.hy926 cells; red bold: SCRB1-overexpressing EA.hy926 cells; red italics: SCRB1-silenced EA.hy926 cells; blue: HAECs; green: HEPG2 cells.

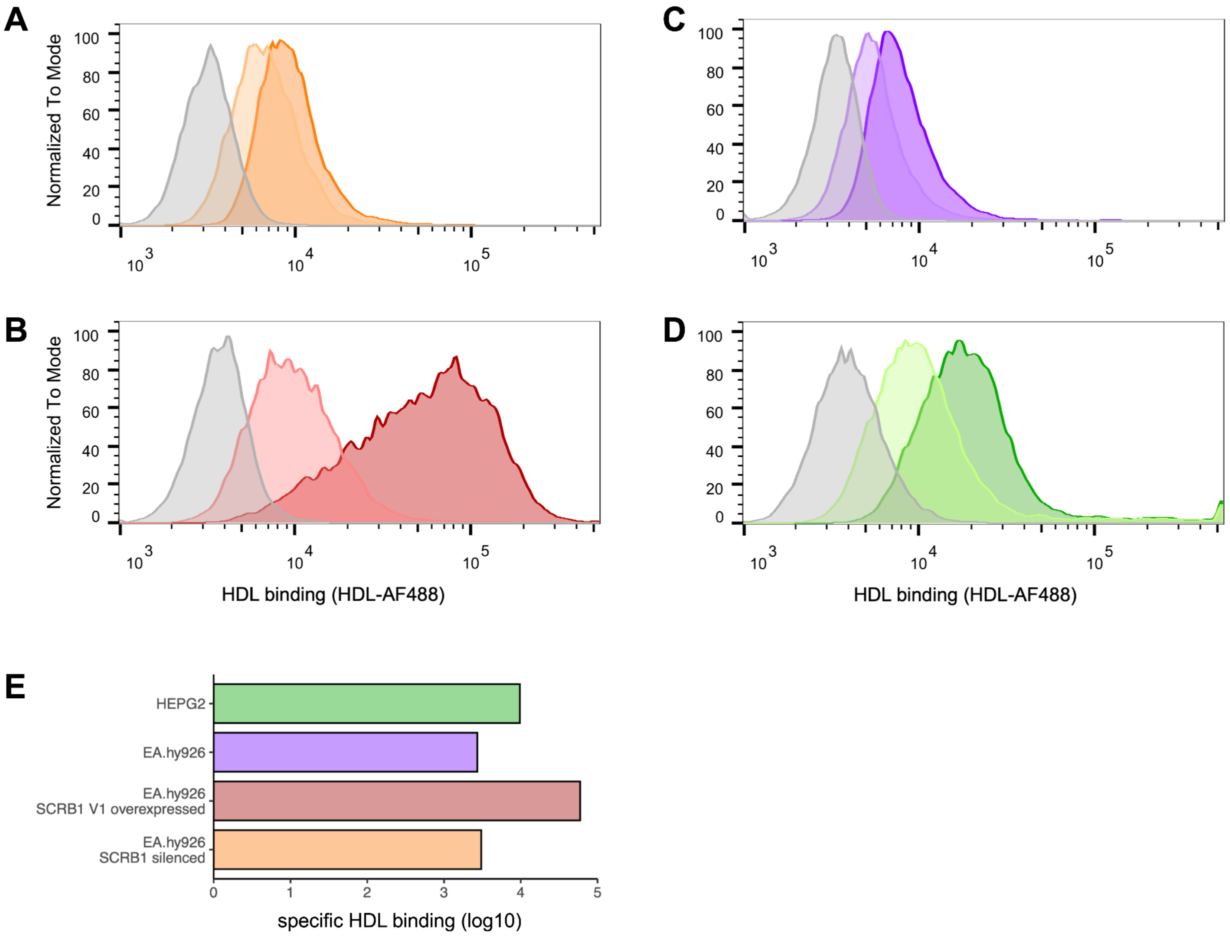


**Suppl. Figure 3:** Flow cytometry-based analysis of HDL binding to various cell types. **A-D)** Flow cytometry analyses of A) SCRB1-silenced EA.hy926 cells, B) SCRB1-overexpressing EA.hy926 cells, C) wild-type EA.hy926 cells, and D) HEPG2 cells. Gray peaks correspond to the untreated controls, dark-colored peaks represent cells incubated with HDL-AF488 in the absence of unlabeled HDL (total binding), light-colored peaks represent cells incubated with HDL-AF488 in the presence of 100-fold excess of unlabeled HDL (non-specific binding). **E)** Specific HDL binding to indicated cell lines, calculated as the difference between total and non-specific binding.

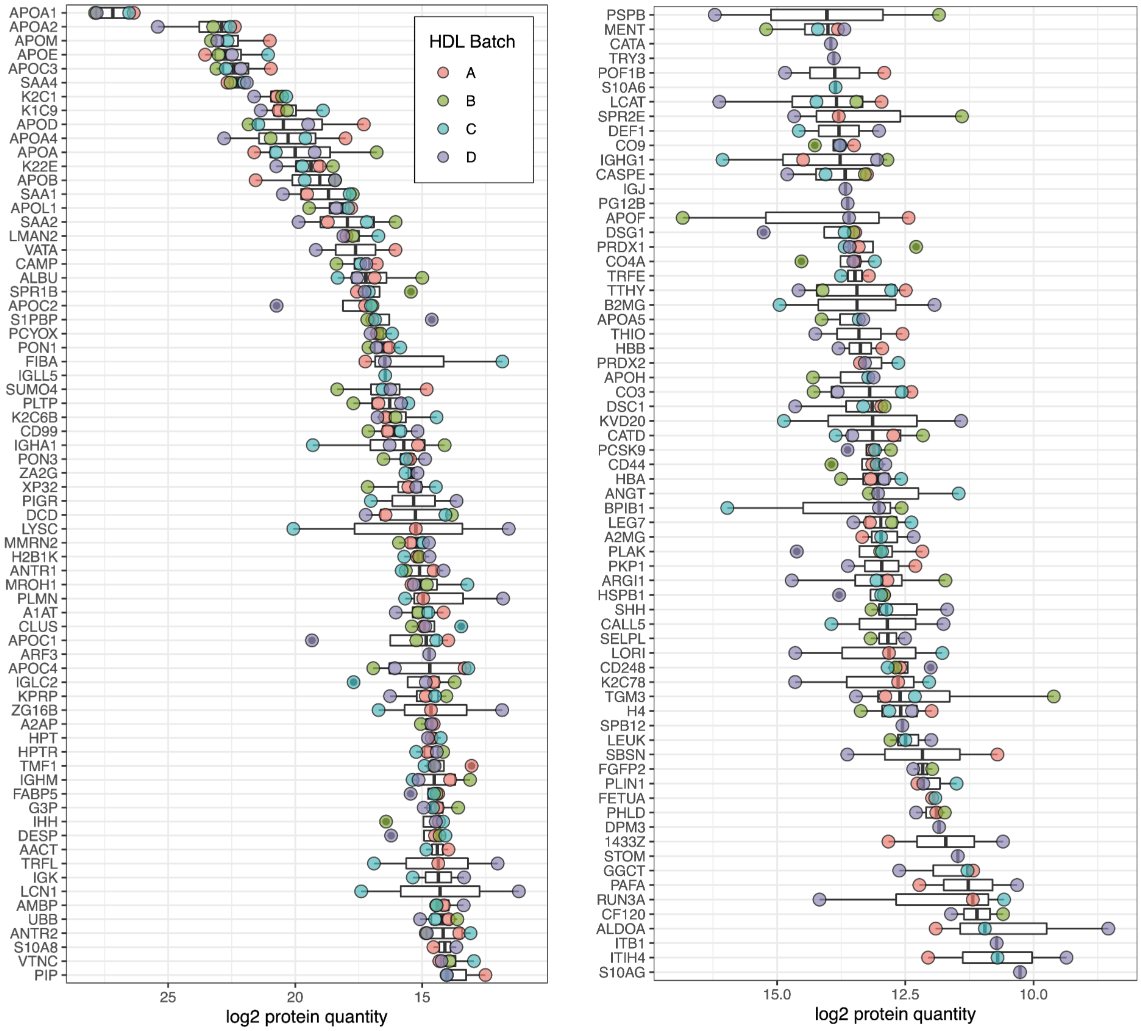


**Suppl. Figure 4:** Protein abundance ranking plots in independent LUX-MS snapshots obtained using different batches of HDLs as ligands on wild-type EA.hy926 cells. The proteins are ordered according to their median abundance across all four HDL isolates.

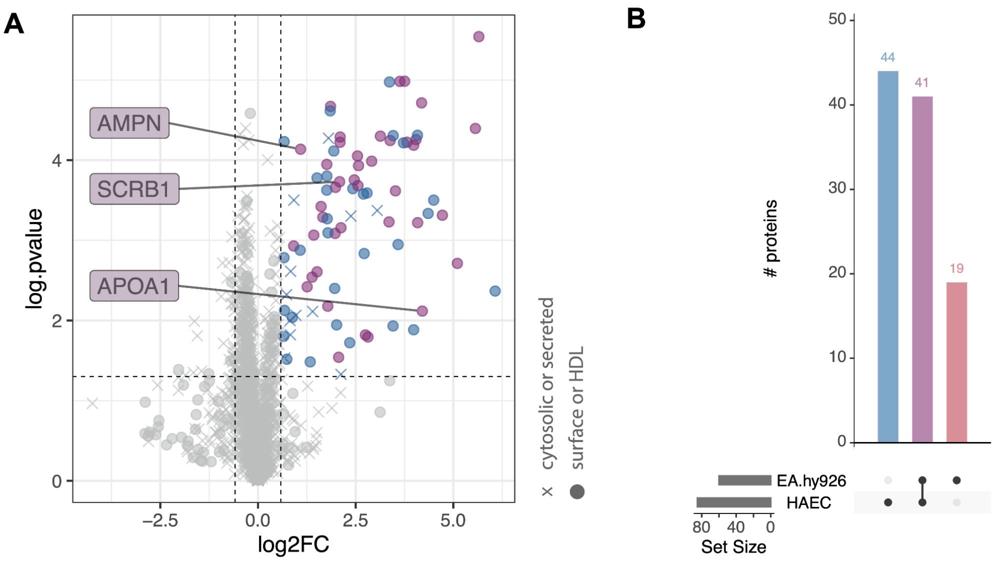


**Suppl. Figure 5:** Validation of the endothelial HDL synapse on HAECs. **A)** Volcano plot of an HDL-guided LUX-MS experiment on HAECs. Blue: Proteins enriched on HAECs. Purple: Proteins of the EA.hy926 HDL synapse core candidate list. **B)** Number of enriched proteins in EA.hy926 cells (blue) and HAECs (red) and the overlap between HAECs and HDL synapse core candidates (purple). Significance was determined using MSstats (FC > 1.5 and p value < 0.05).

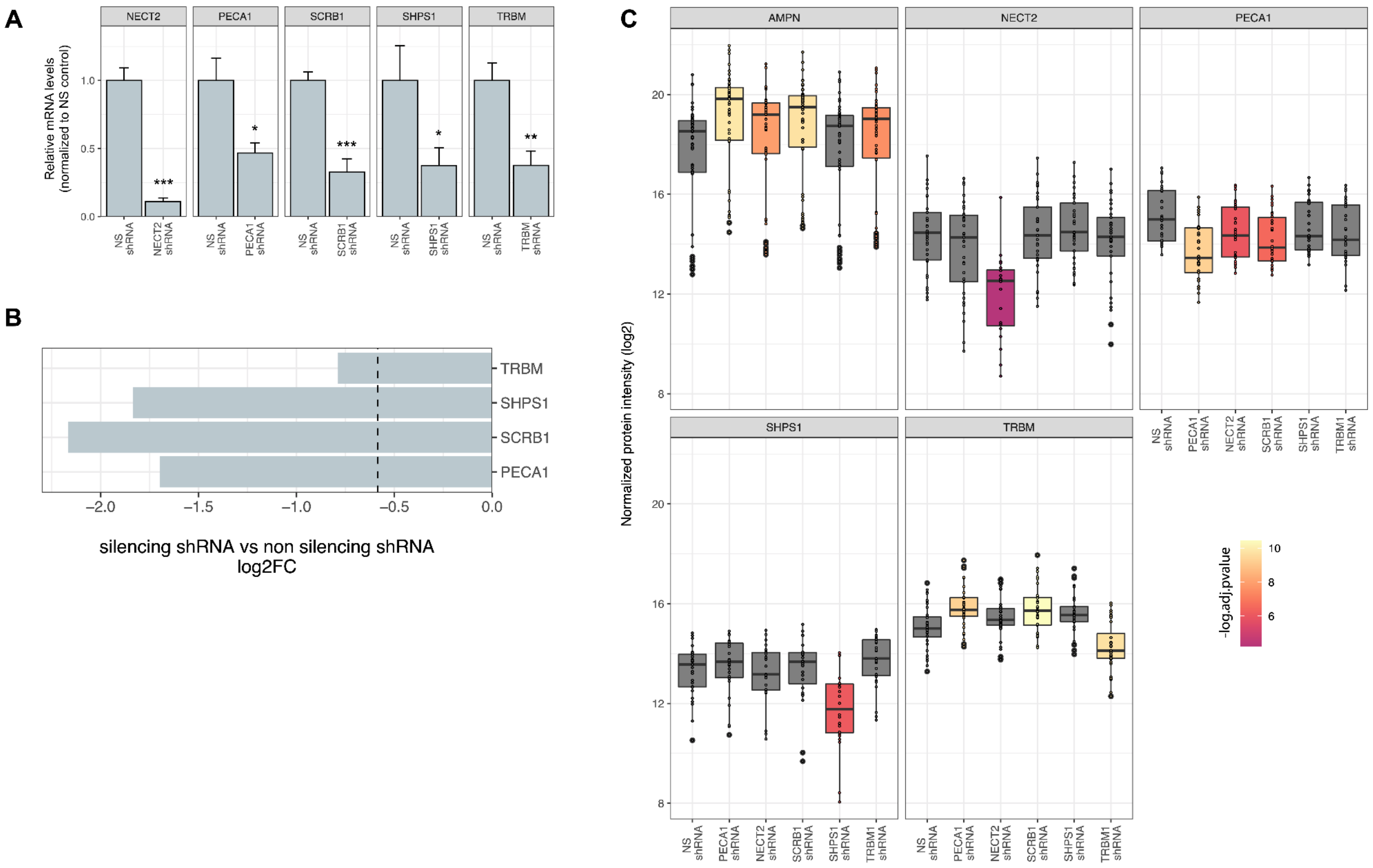
A
NECT2
PECA1
SCRB1
SHPS1
TRBM
Relative mRNA levels (normalized to NS control)
1.0
0.5
0.0
***
*
***
*
**
NS shRNA
NECT2 shRNA
PECA1 shRNA
SCRB1 shRNA
SHPS1 shRNA
TRBM shRNA
B
TRBM
SHPS1
SCRB1
PECA1
−2.0
−1.5
−1.0
−0.5
0.0
silencing shRNA vs non silencing shRNA log2FC
C
AMPN
NECT2
PECA1
SHPS1
TRBM
Normalized protein intensity (log2)
20
16
12
8
NS shRNA
PECA1 shRNA
NECT2 shRNA
SCRB1 shRNA
SHPS1 shRNA
TRBM1 shRNA
-log.adj.pvalue
10
8
6

**Suppl. Figure 6:** Evaluation of silencing efficiencies in EA.hy926 cells. **A)** mRNA abundance levels in cells that express indicated shRNAs normalized to the levels in cells that express control shRNA (NS). The data are shown as means ± SD, and significance was determined with the Student's t-test (* $p < 0.05$; ** $p < 0.01$; *** $p < 0.001$). **B)** Effect of shRNA expression on surface abundance of indicated proteins as determined by auto-CSC. Bars correspond to FC in protein expression in cells that express indicated shRNA compared to control cell line. The dashed line indicates a FC threshold of log2(1.5). NECT2 was not detected on the control cell line. **C)** Effect of shRNA expression on the abundance of candidate proteins in cell lysates determined by mass spectrometry. The color grade represents adj. p value, and gray boxes are not significant. Significance threshold: FC > 1.5 or < -1.5 and adj. p value < 0.05. AMPN protein abundance levels were included. SCRB1 was not detected. Vertical lines indicate minimum and maximum values; boxes indicate first and third quartiles; and horizontal lines indicate medians. Black dots represent single peptide measurements.

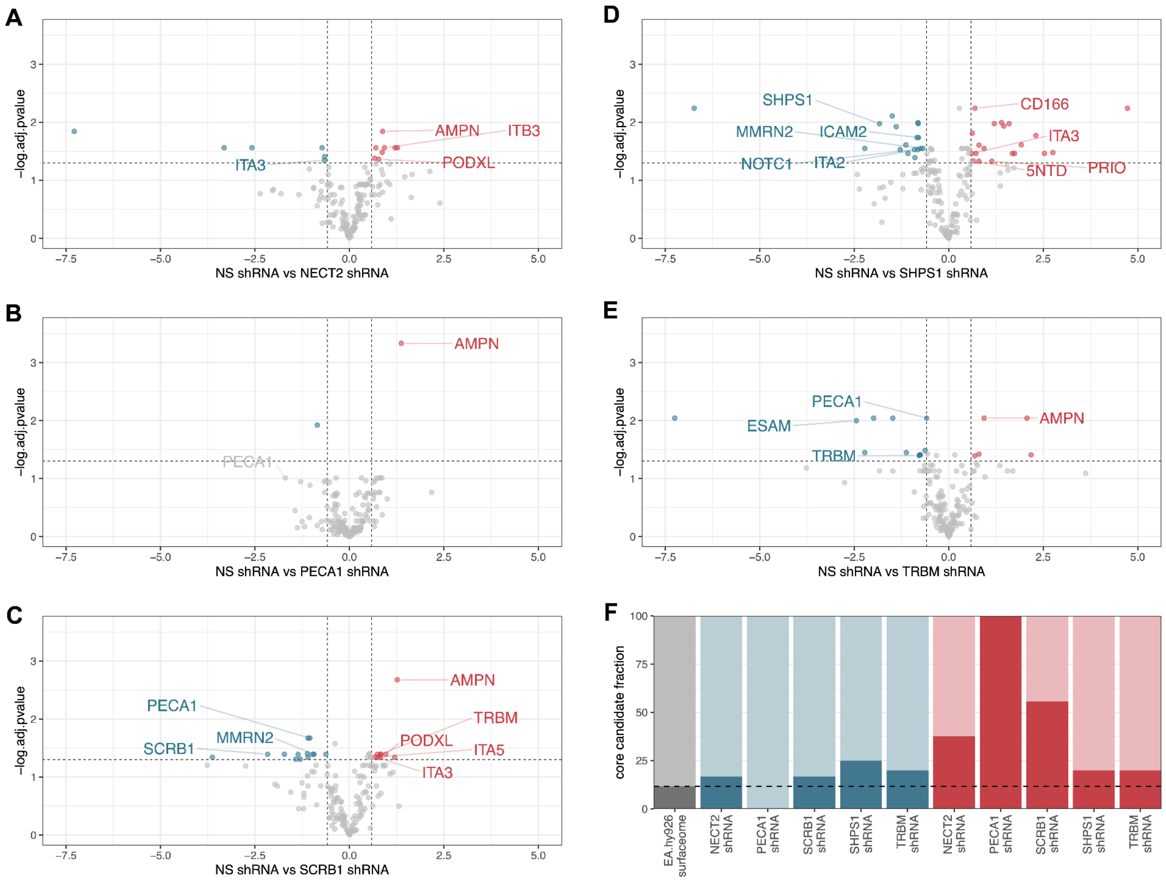


**Suppl. Figure 7:** Quantitative surfaceome analysis of EA.hy926 cells upon shRNA-mediated silencing of **A)** NECT2, **B)** PECA1, **C)** SCRB1, **D)** SHPS1, or **E)** TRBM. Highlighted are significant proteins that belong to the group of 60 HDL synapse core candidates. Significance threshold: FC > 1.5 or < -1.5 and adj. p value < 0.05. **F)** Fraction (%) of up- or downregulated core proteins compared to the total number of upregulated (red) or downregulated (blue) proteins. The shRNA-targeted proteins were excluded from the corresponding groups. The gray bar and the dashed line represent the expected fraction of core candidates (i.e., the fraction of auto-CSC detectable core candidates compared to the total number of EA.hy926 surfaceome proteins established with auto-CSC).

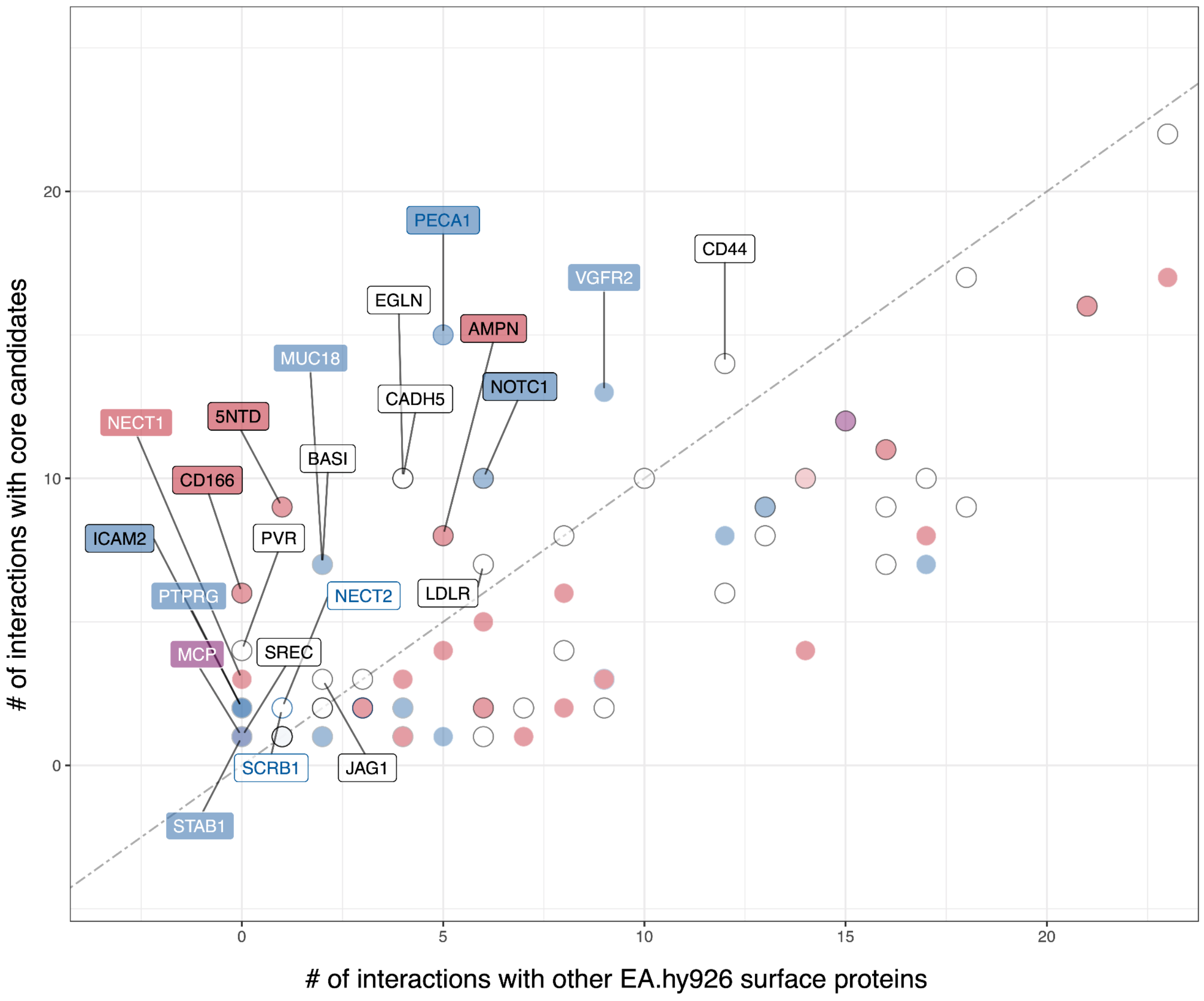


**Suppl. Figure 8:** Comparison of the number of interactions of regulated or core candidates with core candidates against the number of interactions with other EA.hy926 surfaceome proteins. Interactions were extracted from the STRING database with a confidence > 0.7. Highlighted are proteins with more interactions to core candidates than with other EA.hy926 surface proteins. Blue labels indicate proteins downregulated in the auto-CSC experiments with shRNA silenced candidate genes. Red labels indicate proteins upregulated in the auto-CSC experiments with shRNA silenced candidate genes. Black frame indicates core candidate proteins. Blue frame indicates gene that was silenced.

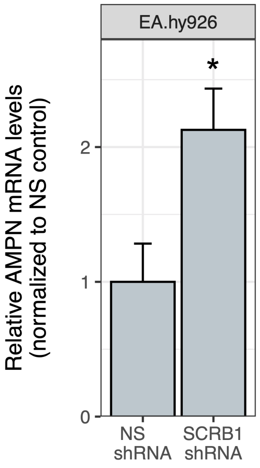


**Suppl. Figure 9:** AMPN mRNA levels in EA.hy926 cells that express shRNA targeting SCRB1 and in cells that express control shRNA (NS). The data are shown as means ± SD, and significance was determined with the Student's t-test (* $p < 0.05$).

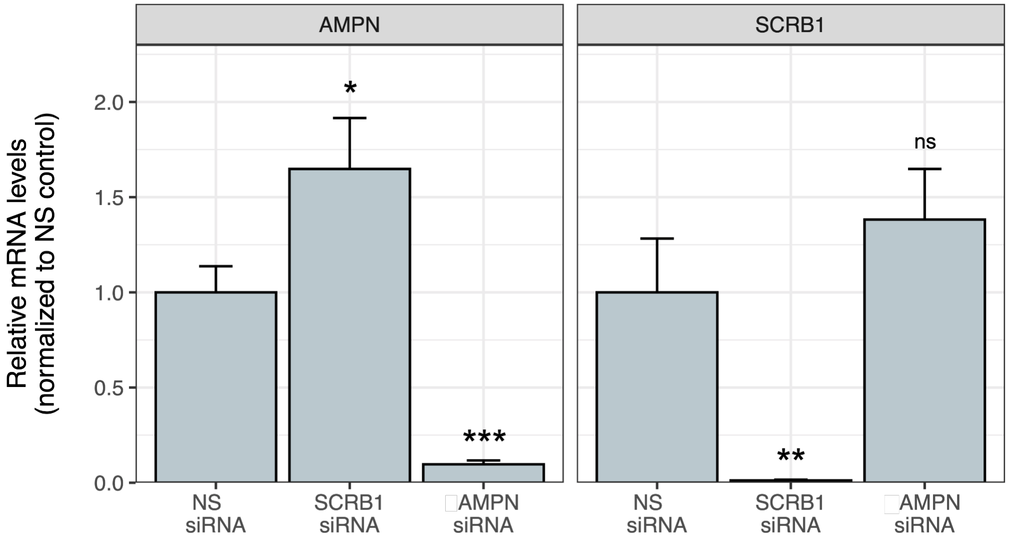


**Suppl. Figure 10:** Relative abundances of AMPN and SCRB1 mRNAs in HAECs upon treatment of cells with indicated siRNAs. Values were normalized to cells treated with control siRNA (NS). The data are shown as means ± SD, and significance was determined with the Student's t-test (* $p < 0.05$; ** $p < 0.01$; *** $p < 0.001$; ns = not significant).

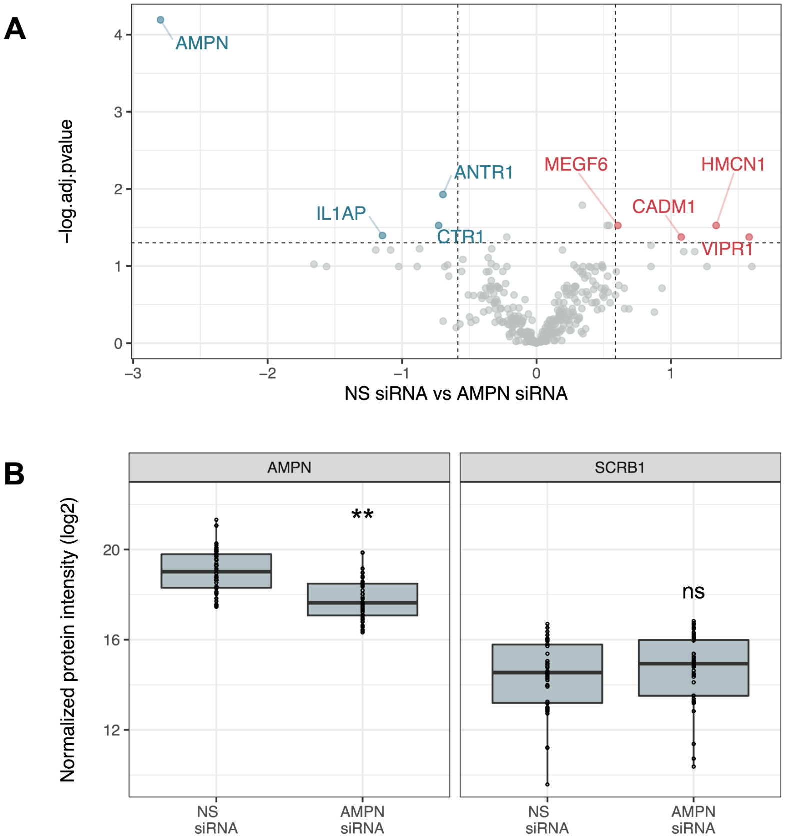


**Suppl. Figure 11:** HAEC surfaceome dynamics upon AMPN silencing. **A)** Quantitative HAEC surfaceome analysis upon AMPN silencing. Highlighted are all significantly upregulated (red) and downregulated (blue) proteins (FC > 1.5 or < -1.5 and adj. p value < 0.05). **B)** Abundances of AMPN and SCRB1 proteins in lysates of HAECs treated with siRNA targeting AMPN compared to the control. Vertical lines indicate minimum and maximum values, boxes indicate first and third quartiles, and horizontal lines indicate median. Black dots represent single peptide measurements. Significance was determined using MSstats (** adj. p value < 0.01; ns = not significant).
